# Drivers of methane-cycling archaeal abundances, community structure, and catabolic pathways in continental margin sediments

**DOI:** 10.1101/2024.11.29.625990

**Authors:** Longhui Deng, Damian Bölsterli, Clemens Glombitza, Bo Barker Jørgensen, Hans Røy, Mark Alexander Lever

**Author notes:** **Correspondence:** Mark Alexander Lever.

## Abstract

Marine sediments contain Earth’s largest reservoir of methane, with most of this methane being produced and consumed *in situ* by methane-cycling archaea. While numerous studies have investigated communities of methane-cycling archaea in hydrocarbon seeps and sulfate-methane transition zones, little is known about how these archaea change from the seafloor downward in the far more common diffusion-dominated marine sediments. Focusing on four continental margin sites of the North Sea-Baltic Sea transition, we here investigate the in *situ* drivers of methane-cycling archaeal community structure and metabolism based on geochemical and stable carbon-isotopic gradients, functional gene (*mcr*A) copy numbers and phylogenetic compositions, as well as thermodynamic calculations. We observe major vertical and lateral changes in community structure that largely follow changes in organic matter reactivity and content, sulfate concentration, and bioturbation activity. While methane-cycling archaeal communities in bioturbation and sulfate reduction zones are dominated by known methyl-dismutating taxa within the *Methanosarcinaceae* and putatively CO_2_-reducing *Methanomicrobiaceae*, the communities change toward dominance of known methane-oxidizing taxa (ANME-2a-b, ANME-2c, ANME-1a-b) in sulfate-methane transitions. Underlying methanogenesis zones were characterized by a change toward mainly physiologically uncharacterized groups, including ANME-1d and several new genus-level groups of putatively CO_2_-reducing *Methanomicrobiaceae* and methyl-reducing *Methanomassiliicoccales*. Notably, group-specific increases in mcrA copy numbers by 2 to 4 orders of magnitude from the sulfate reduction zone into the sulfate-methane transitions or methanogenesis zones indicate the thriving of several major methane-cycling archaeal taxa. Together our study provides insights into the community and pathway shifts vertically along the geochemical gradients and horizontally along the different sedimentary settings and their underlying drivers in continental margin sediments.

## Introduction

Despite being Earth’s largest methane reservoir, marine sediments are only minor sources of atmospheric methane compared to freshwater sediments (Reeburgh, 2007). The high concentration of sulfate in seawater restricts most microbial methane production to deeper sediment layers beneath the sulfatic zone and generally promotes the anaerobic oxidation of >90% of marine sedimentary methane before it can reach the seafloor or overlying water (Reeburgh, 2007). Nonetheless, recent studies suggest that methane release from marine sediments is higher than previously thought, at least in coastal and continental shelf environments (Weber et al., 2019; Lapham et al., 2024). Marine methane emissions may, moreover, increase in the future due to eutrophication and climatic warming (James et al., 2016). The latter promote water column stratification and bottom water oxygen depletion and thereby negatively impact the populations and activities of sediment macrofauna, which drive much of the sedimentary sulfate input (Bianchi et al., 2021; Dean et al., 2018).

Most of marine sedimentary methane is internally produced and consumed by methane-cycling archaea. Methanogens produce methane by converting microbial fermentation products, such as H_2_, acetate, or methanol, to methane via a process known as methanogenesis (Schink, 1997). The distribution of methanogens is partially controlled by competition with respiring microorganisms. Organisms that use oxygen, nitrate, metal oxides (Mn(IV), Fe(III)), or sulfate as electron acceptors for respiration typically have higher energy gains from the same energy substrates than methanogens (Lovley & Goodwin, 1988). As a result, methanogenesis often only dominates respiration in deeper sediments (methanogenesis zones) where these energetically superior electron acceptors are depleted (Jørgensen & Kasten, 2006). A major fraction of this methane diffuses into sediments overlying methanogenesis zones, so-called sulfate-methane transitions (SMTs), where it is consumed by the anaerobic oxidation of methane (AOM) involving close relatives of methanogenic Archaea. These anaerobic methanotrophic Archaea (ANME), in many cases through syntrophic associations with bacteria, couple AOM to the reduction of sulfate (Boetius et al., 2000), metal oxides (Ettwig et al., 2016), or nitrate (Haroon et al., 2013). AOM coupled to sulfate reduction is by far the most important methanotrophic pathway in anoxic marine sediments (Egger et al., 2018).

Multiple archaeal taxa have been linked to methanogenesis and AOM in marine sediments. The dominant methanogens belong to the orders *Methanomicrobiales*, -*sarcinales*, -*cellales*, and -*bacteriales*, all of which are members of the phylum *Euryarchaeota* (Lever, 2013; Wen et al., 2017). In addition, several new groups have been discovered over the last decade. These include cultured methanogenic isolates of the euryarchaeotal order *Methanomassiliicoccales* (Borrel et al., 2013; Dridi et al. 2012) and euryarchaeotal class *Methanonatronarchaeia* (Sorokin et al., 2018), as well as recently members of the phyla Thermoproteota (formerly Verstraetearchaeota; Kohtz et al., 2024) and Korarchaeota (Krukenberg et al., 2024).

Known ANMEs (all Euryarchaeota) are frequently divided into three distinct clades (ANME-1, -2, and -3), based on 16S rRNA genes and genomic phylogenies. ANME-1 (“Candidatus Methanophagales”) is an order-level clade, ANME-2 and ANME-3 belong to the order Methanosarcinales. ANME-2 can be subdivided into four clades: genus-level ANME-2a (“Candidatus Methanocomedenaceae”), genus-level ANME-2b (“Candidatus Methanomarinus”), family-level ANME-2c (“Candidatus Methanogasteraceae”), and family-level ANME-2d (Methanoperedenaceae), whereas ANME-3 (“Candidatus Methanovorans”) represents a genus within the family Methanosarcinaceae (Chadwick et al., 2022). All three ANME clades commonly co-occur in marine cold seep and mud volcano sediments, where they can be among the dominant groups of microorganisms (Lloyd et al., 2010; Takano et al., 2018). By contrast, in diffusion-dominated sediments, ANMEs only account for a small minority of the microbial population and are typically dominated by ANME-1. In these sediments, ANME-1 are frequently found not only in the SMT but also in underlying methanogenesis zone (Beulig et al., 2019; Lever et al., 2023; Yanagawa et al., 2011).

All known archaeal methanogens produce methane via the reductive acetyl CoA pathway, and reduce methyl coenzyme M to methane via methyl coenzyme M reductase as a terminal step (Liu & Whitman, 2008). Four different variations of this pathway are known, which differ in carbon substrates: a) CO_2_ reduction involving H_2_ and formate (‘hydrogenotrophic’) and in exceptional cases primary and secondary alcohols or carbon monoxide; b) acetate disproportionation (‘aceticlastic’); c) dismutation of methylated compounds, e.g. methanol, methyl amines, or methyl sulfides, with or without H_2_ (‘methylotrophic’; all reviewed in Whitman et al. 2014); and d) O-demethylation of methoxylated aromatic compounds (methoxydotrophic; Mayumi et al., 2016). In addition, some methanogens, e.g., *Methanothrix* and *Methanosarcina* (both *Methanosarcinales*), perform CO_2_ reduction by extracellular electron transfer (EET) via conductive structures, e.g. cytochromes, or perhaps pili (Gao & Lu, 2021a; Rotaru et al., 2014) that connect to partner organisms, minerals or organic carbon compounds. Most biogenic methane is believed to be produced via the aceticlastic and hydrogenotrophic reactions (Conrad, 1999; Conrad, 2020), with stable carbon isotopic analyses indicating that CO_2_ reduction prevails in methanogenesis zones of marine sediments (Whiticar, 1999; Whiticar et al., 1986; Beulig et al. 2018). This inference is mainly based on measurements indicating that CO_2_ reduction produces more negative carbon isotopic signatures (δ^13^C-CH_4_: -60 to -110 ‰) than aceticlastic methanogenesis (δ^13^C-CH_4_: -50 to -60 ‰). By contrast, methylotrophic methanogenesis has been shown to occur and dominate methanogenic processes in sulfate-rich marine surface sediments affected by bioturbation (Xiao et al., 2018; Zhuang et al., 2018). This production of methane from methylated compounds has been attributed to these compounds not being used by most sulfate reducers (“non-competitive substrates”; (King, 1984; Oremland et al., 1982).

To date, most research on sedimentary methane-cycling archaea has focused on advective systems, such as hydrocarbon seeps (Knittel et al., 2005; Lloyd et al., 2010; Niu et al., 2022; Orcutt et al., 2010; Ruff et al., 2018; Takano et al., 2018). In addition, sulfate-methane transitions of diffusion-dominated marine sediments have been a major research focus (Beulig et al., 2019; Harrison et al., 2009), as anaerobic methane-oxidizing archaea inhabiting SMTs almost fully consume the highly potent greenhouse gas methane and thus prevent it from escaping to overlying water and to the atmosphere. In contrast, much less is known about the community structure and pathways of methanogenesis and AOM in bioturbated surface sediments (BTZ), sulfate reduction zones (SRZ), or methanogenesis zones (MGZ) of diffusion-dominated sediments, and how these communities and pathways change vertically in response to different geochemical conditions or horizontally to different sedimentary settings.

In this study we explore the diversity, community structure and pathways of methane-cycling communities and their potential environmental drivers at four continental margin sites at the North Sea-Baltic Sea transition. We integrate geochemical, stable isotopic, Gibbs energetic, and methane-cycling archaeal abundance and community data across sites that range from coastal eutrophic to off-shore oligotrophic and differ greatly in sedimentation rates, organic carbon inputs, electron acceptor distributions, as well as microbial and macrofaunal activity and community structure (Deng et al., 2020). Sediment cores from three of the locations extend well into the methanogenesis zone and were sampled at high depth resolution across the SMTs, thus allowing for detailed analyses of methanogenic and methanotrophic community shifts across this important biogeochemical transition interval. Based on our comprehensive geochemical and microbiological data, we identify key drivers of methanogenic and methanotrophic communities in continental margin sediments.

## Materials and methods

### Site description

All samples were taken during a cruise of the R/V *Aurora* in August-September 2014. The four sites AU1 (586 m water depth), AU2 (319 m), AU3 (43 m), and AU4 (37 m) are located along a water-depth gradient between the North Sea and Baltic Sea (**Figure 1**). Sedimentation rates decrease with increasing water depth and distance to shore, and are 0.14, 0.27, 0.30, and 0.33 cm yr^-1^ at AU1-4, respectively (Deng et al., 2020). AU1 and AU2 are in the Skagerrak region, with AU1 being located near the bottom of the Norwegian Trench and AU2 on the southern slope of the same trench. Both sites receive silty clay with low-reactivity allochthonous OM through lateral transport and have high rates of iron and manganese reduction in the top 10 cm (Kristensen et al., 2018). AU3 in the northern Kattegat is dominated by fine sands, while AU4 in the Lillebælt region of the Baltic Sea is dominated by silty clay. AU4 is subject to seasonal bottom water hypoxia and is sulfidic with the exception of a ∼1-mm-thick oxidized surface layer. Macrofaunal biomass increases from AU1 to AU3, but macrofauna is absent from AU4 (Deng et al., 2020; Kristensen et al., 2018).

**Figure 1.**
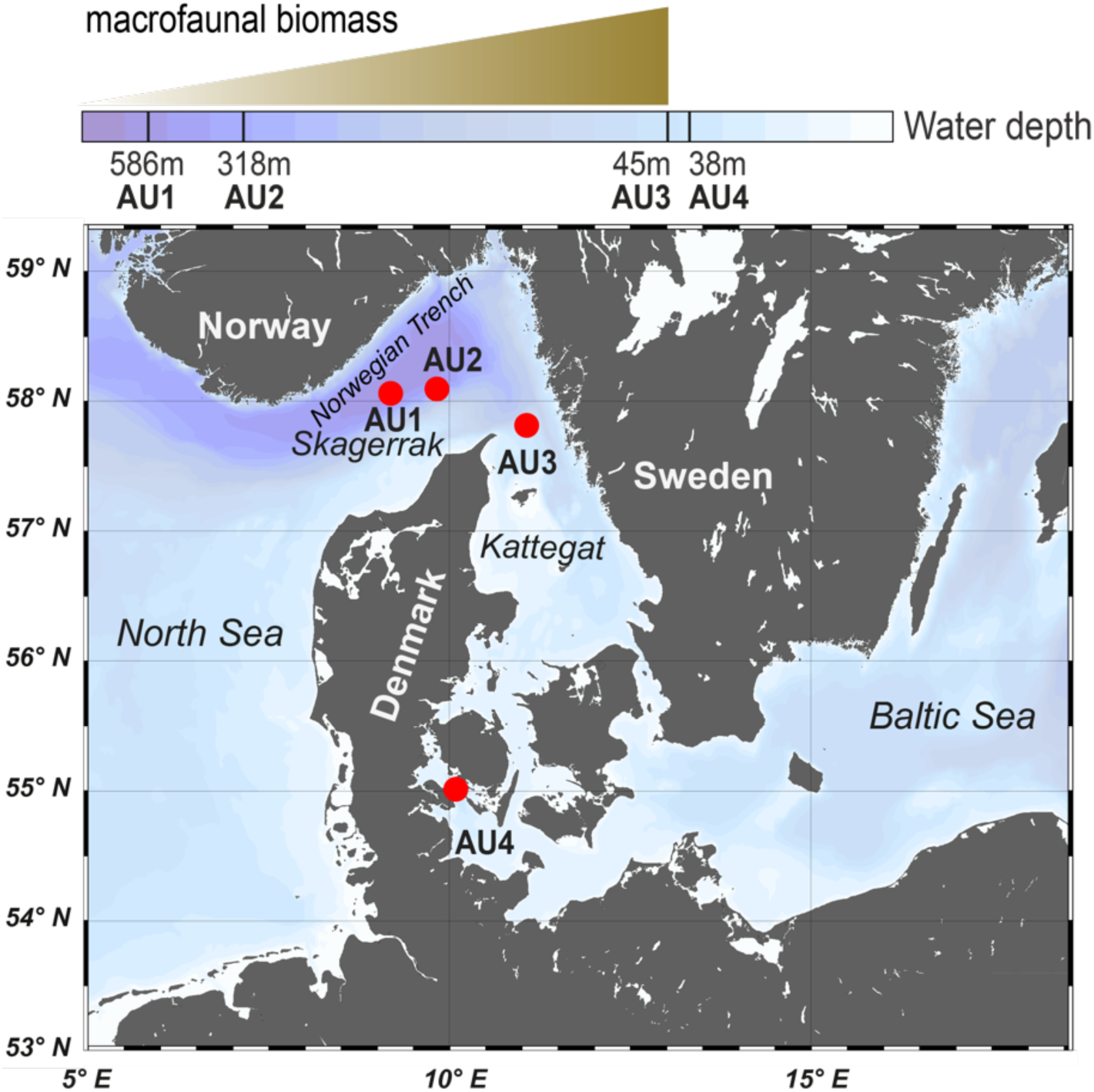
Map of the Skagerrak-Kattegat-Belt Sea area. Four sampling sites (AU1-AU4) are indicated by red dots, along a decreasing gradient of water depth. Macrofaunal biomass increases from AU1 to AU3, while no bioturbation was detected at AU4.

### Sampling scheme

The top 40-50 cm of sediments were sampled using a Rumohr corer, a lightweight corer that enables recovery of nearly undisturbed surface sediments. All deeper sediments (to 500 cm below seafloor (cmbsf)) were collected using a gravity corer. Sediment porewater was sampled from Rumohr cores in 5-cm intervals and from Gravity cores in 10-cm intervals to 100-cm depth followed by 25-cm intervals below. Porewater was extracted using rhizon samplers (Rhizosphere, The Netherlands) that were inserted horizontally through holes drilled into the side of the PVC core liners. Of the sampled porewater samples, 1 ml was preserved with 10 μl saturated HgCl_2_ at 4°C for DIC and δ^13^C-DIC analyses, 1 ml was stored at 4°C for dissolved SO_42-_ quantification, and 2-4 ml was frozen at -20°C for analyses of volatile fatty acid (VFA) concentrations. Subsequently, the Rumohr cores were extruded in 2-cm intervals to 20-cm depth followed by 4-cm intervals to 48-cm depth, while the gravity cores were sampled at 8-10 cm depth intervals. All sediment for DNA, methane, δ^13^C-methane, total organic carbon (TOC), and δ^13^C-TOC analyses was sampled using 5-ml sterile, cut-off syringes. Samples for DNA, TOC, and δ^13^C-TOC analyses were immediately frozen at -20°C and transferred to -80°C upon arrival at the home laboratory. Samples for analyses of methane concentrations and δ^13^C-methane were preserved in saturated NaCl (6M) and stored at 4°C until measurement.

### DNA extraction

DNA was extracted from sediments following lysis protocol II of Lever et al. (2015). This protocol combines chemical (lysis solution I) and mechanical cell lysis (bead-beating: 0.1-mm Zirconium beads), 2× washing with chloroform:isoamyl alcohol (24:1), and precipitation with linear polyacrylamide, NaCl and ethanol. DNA was purified according to protocol A of the CleanAll DNA/RNA Clean-Up and Concentration Micro Kit (Norgen Biotek Corp., Canada).

### Quantitative PCR (qPCR)

*mcr*A copy numbers in DNA extracts were quantified on a LightCycler 480 II (Roche Life Science, Switzerland) by SYBR-Green based qPCR assays using the Mlas_F (5’-GGT GGT GTM GGD TTC ACM CAR TA -3’) / McrA-rev (5’-CGT TCA TBG CGT AGT TVG GRT AGT -3’) primer pair (Steinberg & Regan, 2009). The thermal cycler protocol consisted of (1) enzyme activation and initial denaturation at 95 °C for 5 min; (2) 50 cycles of denaturation at 95 °C for 10 s, annealing at 53 °C for 20 s, elongation at 72 °C for 30 s, and fluorescence measurement at 84 °C for 5 s; and (3) a stepwise melting curve from 95 °C to 53 °C in 1 min intervals to check for primer specificity. Plasmids of *mcr*A from *Methanocorpusculum parvum* were applied as standards. All standards and samples were measured in duplicate.

### Sequencing and bioinformatics

*mcr*A sequence libraries were prepared according to a published workflow (Deng et al., 2020). Using the same primer pair as for qPCR, *mcr*A amplicons were sequenced using the Illumina MiSeq platform (Illumina Inc., California, USA). Raw reads were quality-checked by *FastQC* (http://www.bioinformatics.babraham.ac.uk/projects/fastqc), read ends trimmed using *seqtk* (https://github.com/lh3/seqtk), paired end reads merged using *flash* (Magoč & Salzberg, 2011), primer sites trimmed by *usearch* (Martin, 2011), and quality filtering done by *prinseq* (Schmieder & Edwards, 2011). Zero-radius Operational Taxonomic Units (ZOTUs) were generated using the UNOISE3 algorithm (Edgar, 2016) and clustered using a 97% identity threshold to generate 97% ZOTUs’ (referred to as ‘ZOTUs’ from now on). Taxonomic assignments were done in ARB (www.arb-home.de) using neighbor-joining phylogenetic trees with Jukes Cantor correction and 1,000 bootstrap replicate calculations. These trees were based on an up-to-date, public database (*mcrA4All*; https://drive.google.com/drive/folders/1G8GeJuYsIX4MLv5-LaUQHD9f9F9rfIAu; (Lever et al., 2023) with >2,400 high-quality, optimally aligned *mcr*A sequences from pure culture, amplicon sequencing, metagenome, and whole-genome studies.

### Geochemical analyses

Depth profiles of TOC and 8^13^C-TOC and porewater concentrations of sulfate, methane, and DIC were published previously (Deng et al., 2020; Marshall et al., 2019). The isotopic composition of carbon in DIC and methane were measured on a Thermo Fisher Scientific DeltaV™ isotope ratio mass spectrometer equipped with GasBench II™ for DIC-analysis after liberation via phosphoric acid, and a Precon™ trace gas pre-concentrator for separation and combustion of methane. The δ^13^ C values are reported vs. the Vienna Pee Dee Belemnite standard. Concentrations of volatile fatty acids (VFAs) were determined by 2D-ion chromatography as previously published (Glombitza et al., 2014). The *in situ* pH was calculated from measured alkalinity values (measured by titration) and DIC concentrations.

### Gibbs energy calculations

Gibbs energies (Δ*G_r_*) of (1) methanogenesis reactions from H_2_+CO_2_ (HCO_3_^-^ + 4 H_2_ + H^+^➔ CH_4_ + 3 H_2_O), acetate (CH_3_COO^-^ + H_2_O ➔ CH_4_ + HCO_3_^-^), methanol (4 CH_3_OH ➔ 3 CH_4_ + HCO_3_^-^ + H_2_O + H^+^), and methanol+H_2_ (CH_3_OH + H_2_ ➔ CH_4_ +H_2_O), (2) anaerobic formate oxidation (HCOO^-^ + H_2_O ➔ HCO_3_^-^ + H_2_) and anaerobic acetate oxidation (CH_3_COO^-^ + 4 H_2_O ➔ 2 HCO_3_^-^ + 4 H_2_ + H^+^), and (3) sulfate-dependent AOM (SO_4_^2-^ + CH_4_ ➔ HS^-^ + HCO_3_^-^ + H_2_O) were calculated based on the equation

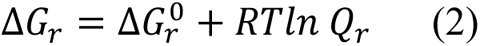

where ΔG*_r_*^0^ is the Gibbs energy (kJ mol^-1^ of reaction) at standard concentrations (1M per each reactant and product, pH 7.0), corrected for *in situ* temperature T (K) and pressure *p* (bar) based on standard enthalpies and molar volumes as outlined in Stumm and Morgan (1996), R is the universal gas constant (0.008314 kJ mol^−1^ K^−1^), and *Q_r_* the quotient of product and reactant activities. Calculations were done for measured concentrations of DIC, acetate, methane, and sulfate, and pH values that were calculated from measured alkalinities and DIC concentrations. For H_2_, methanol, and hydrogen sulfide (HS^-^) concentrations, which were not measured, we performed calculations for concentrations that were assumed to be at the lower and upper *in situ* extremes of these chemicals (H_2_: 0.1 nM and 10 nM; methanol: 1 nM and 1 µM; HS^-^: 1 nM and 10 mM). Calculations involving assumed concentration minima and maxima were used to assess the energetic feasibility of hydrogenotrophic and methylotrophic methanogenesis reactions and AOM, and the conditions or locations where these reactions were most likely to take place. Activities of all aqueous species were calculated from measured concentrations multiplied by their activity coefficients. These were γ_HCO32−_ = 0.532 (Millero et al., 1982), γ_CH4_ = 1.24 (Millero, 2000), γ_SO42−_ = 0.104 (Millero et al., 1982), and γ_HS−_ = 0.685 (Clegg & Whitfield, 1991). The activity coefficient of H_2_ was set to 1.0116, while we assumed activity coefficients of 1.0 for acetate and methanol. Following convention, the activity of H^+^ was equal to its pH-value (e.g. pH 8.0 ➔ 10^-8^), and the activity of H_2_O was set to 1.0. Standard Gibbs energies (**ΔG_f_°)**, standard enthalpies (**ΔH_f_°)**, and standard molal volumes (**ΔV_f_°)** of formation are shown in **Supplementary Table 1**.

### Multivariate statistics

All statistical analyses were performed in R (http://www.R-project.org). Richness (the observed number of ZOTUs), evenness, and Non-metric Multi-Dimensional Scaling (NMDS) based on Bray-Curtis distances of *mcr*A communities between samples were calculated using the ‘phyloseq’ package (McMurdie & Holmes, 2013). PERmutational Multivariate ANalysis Of VAriance (PERMANOVA, permutations = 999) and statistical tests (Welch’s t test and Wilcoxon test) were performed using the “vegan” and “stats” packages, respectively (Oksanen et al., 2013). Correlations between the abundances of *mcr*A groups and environmental variables were calculated and visualized using the “corrplot” package (Wei et al. 2017). All calculations were performed based on ZOTU-level phylogenetic assignments unless stated otherwise.

## Results

### Geochemical profiles related to the sedimentary methane cycle

The four stations show up to 10-fold differences in total organic carbon (TOC) contents, as well as an increase in microbial activity from deep to shallow stations (**Figure 2**).

**Figure 2.**
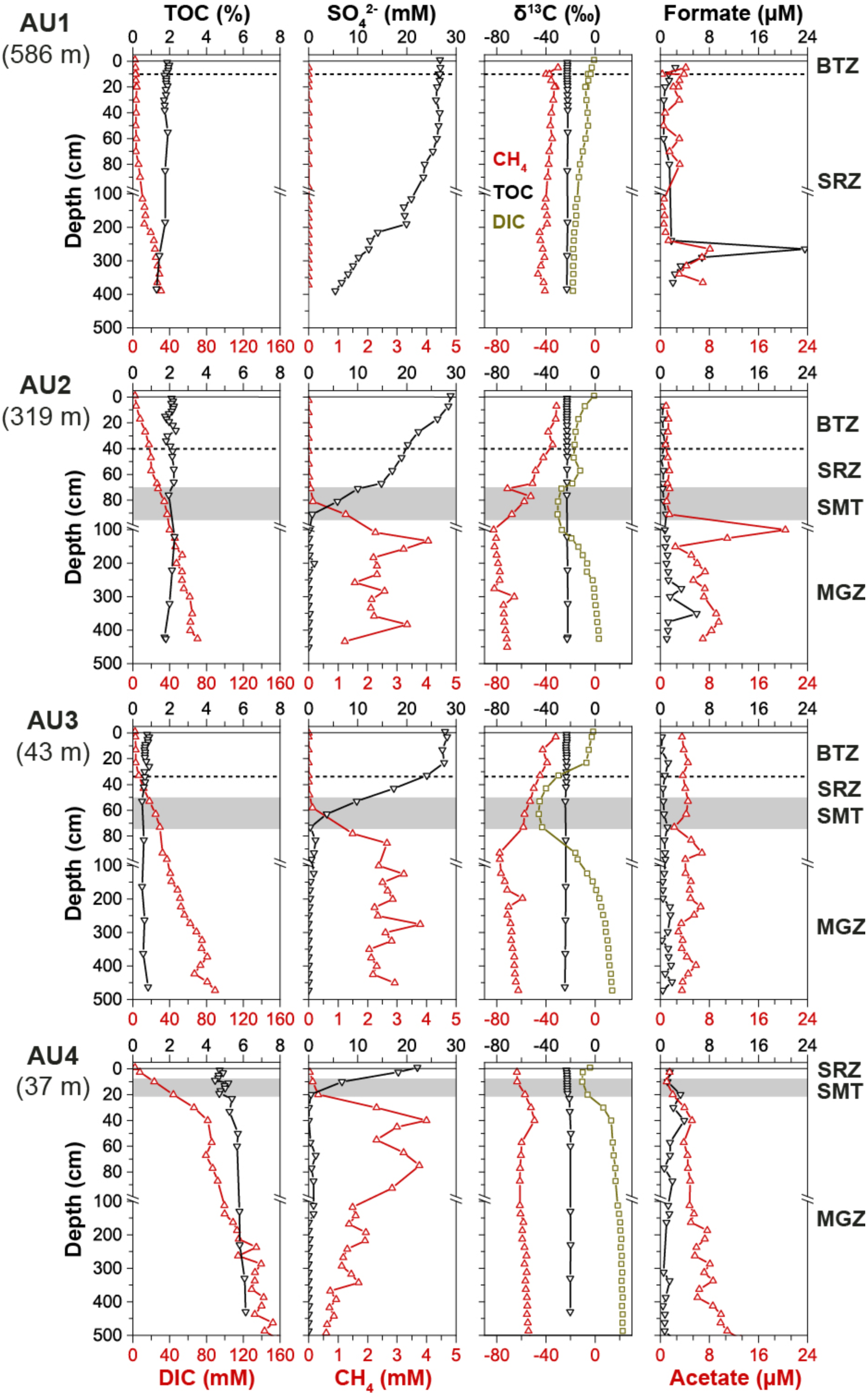
Depth profiles of TOC, DIC, SO_4_^2-^, CH_4_, δ^13^C-TOC, δ^13^C-DIC, δ^13^C-CH4, formate, and acetate at AU1-4. Depth intervals of the bioturbation zone (BTZ), sulfate-reduction zone (SRZ), sulfate-methane-transition (SMT), and methanogenesis zone (MGZ) are indicated by horizontal dashed lines and gray bars, respectively. Note: due to CH_4_ supersaturation and resulting outgassing of CH_4_ at atmospheric pressure, measured CH_4_ concentrations below the SMT are likely to be significant underestimates and do not represent *in situ* concentrations.

TOC contents (% sediment dry wt.) are highest at the sulfidic coastal station (AU4; 4.7-∼6%), lowest at the sandy shallow water station (AU3; ∼0.5-1%), and have intermediate values at the two deep stations (AU1: 1.3-1.9%; AU2; 1.7-2.1%). By contrast, corresponding DIC concentration profiles, used as a proxy for organic matter mineralization rates, indicate increases in DIC concentrations from the deepest (AU1) to the shallowest station (AU4). This apparent increase in mineralization rates from deep to shallow water is also reflected in sulfate concentrations, and most notably the depth of sulfate penetration, which decreases with water depth (AU1: >400 cm, AU2: ∼90 cm, AU3: ∼80 cm, AU4: ∼20 cm). Corresponding methane concentrations remain at background values (≤10 µM) throughout the sulfate-rich AU1 core, but increase to millimolar concentrations in the SMT at the three other stations. SO_4_^2−^ and DIC concentrations are nearly constant in the top 60, 10, and 25 cm of AU1, AU2, and AU3. This is explained by significant bioirrigation activity at these sites, and extensive zones of iron and manganese reduction-dominated microbial respiration at AU1 and AU2 (Deng et al., 2020).

The carbon stable isotopic compositions show clear trends across stations. The δ^13^C-TOC values throughout AU1 to AU3 and in the top 20 cm of AU4 are in the typical range of marine phytoplankton (-24 to -22‰; (Fry & Sherr, 1989). Below 20 cm, the δ^13^C-TOC at AU4 increases slightly (−20‰ at 50 cm and below). The δ^13^C-CH_4_ at AU1 decreases from -30‰ at ∼5 cm to -41‰ at ∼390 cm. At AU2 and AU3, δ^13^C-CH_4_ are also around -30‰ in surface sediments, but decrease throughout the sulfate zone and SMT, reaching their lowest values right below the SMT (-80‰), before increasing slightly in deeper parts of the methanogenesis zone (-70 to -60‰). In marked contrast, at AU4 δ^13^C-CH_4_ increases from -60‰ in surface sediments to -50‰ at ∼40 cm and remains relatively constant below (-58±2‰). The δ^13^C-DIC profiles also show strong variations between sites. At all sites, δ^13^C-DIC values are near seawater values (0‰) in surface sediments. Yet, while at AU1 δ^13^C-DIC values show a gradual decrease with depth to -18‰ at 390 cm, all other stations have unimodal distributions, with the most negative δ^13^C-DIC values in or right above the SMT (AU2: -30‰; AU3: -28‰; AU4: -10‰). In the uppermost section of the methanogenesis zones, the δ^13^C-DIC increases steeply and leads to δ^13^C-DIC > 0‰ below a certain depth in the methanogenesis zone (AU2: ∼3 m; AU3: ∼1.6 m; AU4: ∼0.2 m). δ^13^C-DIC-values at the core bottoms of these stations are at +3‰ (AU2), +14‰ (AU3), and +22‰ (AU4).

Concentrations of formate and acetate are mostly in the low micromolar range (≤10µM) at all four stations, with acetate concentrations exceeding those of formate in most samples. At AU1, both remain below 4 µM from 0-250 cm but show a strong subsurface peak at 265 cm (formate: ∼24 µM, acetate: ∼8 µM), below which concentrations drop again. At AU2, formate and acetate concentrations are <3 µM above the SMT. Below the SMT, formate concentrations increase down to 351 cm (formate: ∼6 µM; acetate: ∼9 µM) before decreasing again towards the core bottom (formate: ∼1 µM; acetate: 7 µM), and acetate shows an additional peak in the uppermost sample of the methanogenesis zone (∼20 µM; 101 cm). By comparison, formate and acetate concentrations are more constant with depth at AU3 (formate: 0.9±0.5 µM, acetate: 4±1 µM). At AU4, both formate and acetate concentrations increase from ∼1.5 µM at 5 cm to 4-5 µM at 40 cm. Below this depth, formate concentrations gradually decrease while acetate concentrations gradually increase (formate: ∼0.7 µM; ∼11 µM).

### Depth-related trends in absolute and relative abundances of mcrA copies

Copy numbers of *mcr*A are similar in surface sediments of all sites but increase from the oligotrophic AU1 to the eutrophic AU4 when deeper layers are compared across these sites. In addition, *mcr*A copies increase across the SMTs into the underlying methanogenesis zones at AU2-4, indicating growth of methane-cycling archaeal populations (**Figure 3A**).

**Figure 3.**
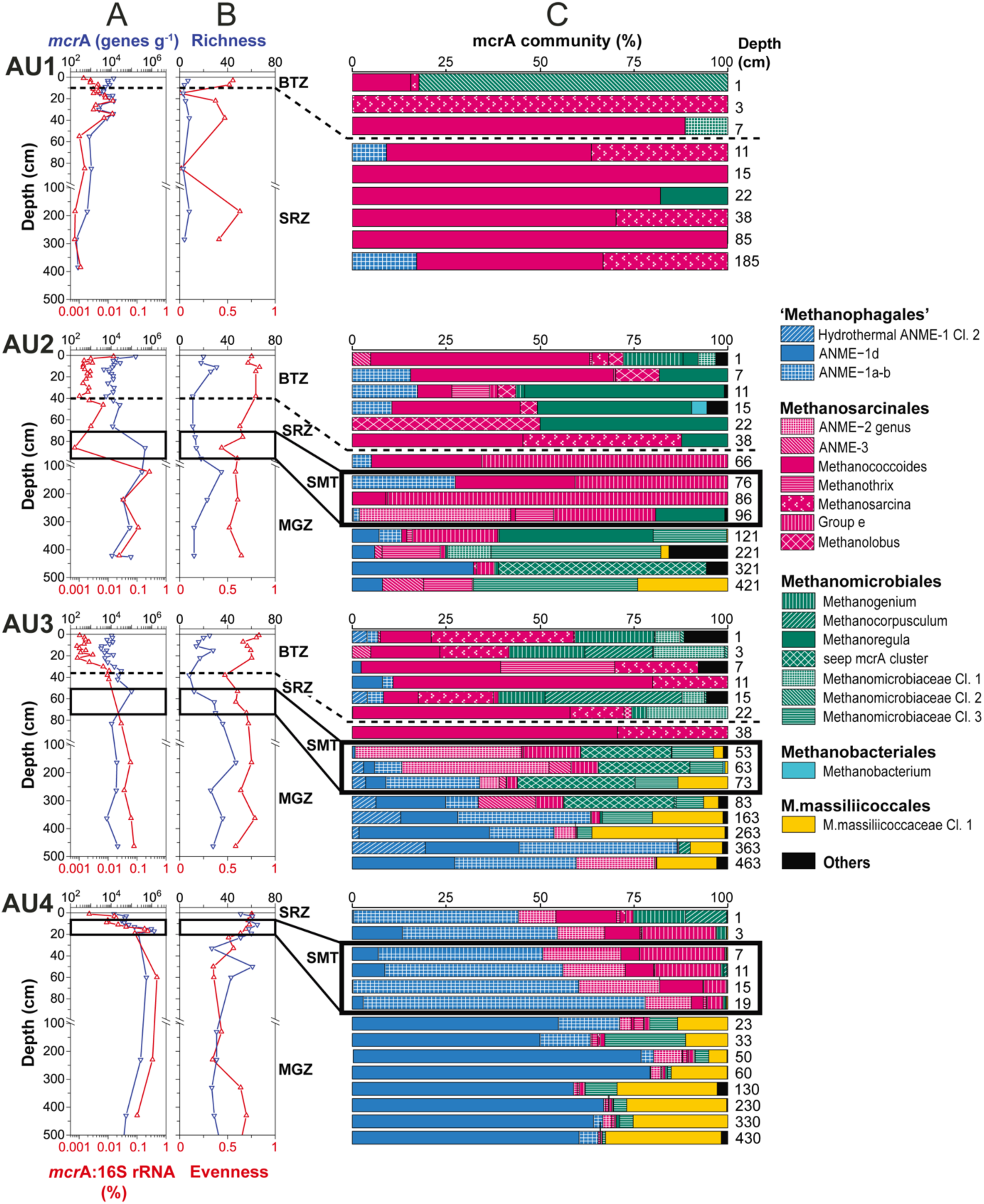
Depth profiles of methane-cycling archaeal communities at AU1-4. (**A**) *mcr*A gene copy numbers and ratios of *mcr*A to total 16S rRNA gene copy numbers; (**B**) richness and evenness based on *mcr*A ZOTUs; (**C**) *mcr*A community composition at the genus-level. Samples above the horizontal dashed lines were located within the bioturbation zone. Black boxes indicate samples that were located within the sulfate-methane transition. [Abbreviations: BTZ = bioturbation zone, SRZ = sulfate-reducing zone, SMT = sulfate-methane transition, MGZ = methanogenic zone].

At AU1, *mcr*A abundances fluctuate around ∼10^4^ gene copies g^-1^ within the upper 40cm and decrease to ∼10^2^ gene copies g^-1^ below. At AU2, except for a local high at the sediment surface (∼10^5^ genes g^-1^), *mcr*A copies are relatively stable (∼10^4^ genes g^-1^) within the bioturbated upper 40 cm, below which they increase to 5×10^5^ copies g^-1^ at and right below the SMT, and then decrease gradually to ∼10^4^ g^-1^ at the core bottom. At AU3, *mcr*A copies are ∼10^4^ g^-1^ in the strongly bioturbated top 20 cm, increase to ∼10^5^ g^-1^ at 53 cm and are relatively stable around 10^4^ g^-1^ below. At AU4, *mcr*A copies increase from ∼10^4^ g^-1^ at the sediment surface to ∼10^6^ g^-1^ around the SMT (∼20 cm) and are relatively constant below (∼10^5^ g^-1^).

Abundances of methane-cycling archaea relative to total prokaryotic communities were estimated based on ratios of *mcr*A to total 16S rRNA gene copy numbers (bacterial+archaeal). These ratios suggest (local) increases in the relative abundances of methane-cycling archaea with sediment depth at all 4 locations. At AU1, which has no methanogenesis zone, these increases are restricted to sediments within or near the bottom of the bioturbation zone (1 cm: ∼0.001%; 22 cm: ∼0.015%), below which they decrease back to ∼0.001% at the core bottom. At the other three stations, relative abundances are in the same range as at AU1 in surface sediment (∼0.001%) and also increase clearly at the bottom of the bioturbation zone (AU2, AU3). However, an additional strong increase occurs further down across the SMT and into the upper layers of the methanogenesis zone, where maximum values of 0.3% (AU2), 0.1% (AU3), and 0.5% (AU4) were detected. While relative abundances of methane-cycling archaea were highest at the core bottom of AU3, relative abundances at AU2 and AU4 decreased again toward the core bottom to values of 0.01% (AU2) and 0.1% (AU4).

### Zonation of major methane-cycling archaeal clades in relation to sites and vertical zones

*mcr*A richness calculated based on ZOTUs increases from AU1 to AU4 (**Figure 3B**). At AU1, richness is low (5±2) at all depths. The other three locations share similar trends, with local peaks in the upper parts of the bioturbation and methanogenesis zone. Notably, richness remains high also in deeper portions of AU3. Community evenness shows high scatter at AU1, depth-related decreases at AU2 and AU4 (though note the increase at the bottom of AU4), and a bimodal distribution, with peaks in the bioturbation and methanogenesis zone, at AU3. Notably, Εvenness is overall highest at AU3.

The community composition of methane-cycling archaea varies greatly across sites and in relation to the zones of bioturbation, sulfate reduction, AOM (SMT), and methanogenesis (**Figure 3C**; for taxonomic assignments see phylogenetic tree in **Figure 4**). Diverse genera of *Methanosarcinales* and *Methanomicrobiales* dominate throughout AU1 and AU2, and down to the SMT of AU3, while the Candidate order Methanophagales (ANME-1) dominates the lower part of the SMT and methanogenesis zone of AU3 and throughout AU4. Notably also, *Methanomassiliicoccales* account for major fractions (∼10-35%) of *mcr*A reads throughout methanogenic sediments of AU3 and AU4 and at the bottom of AU2.

**Figure 4.**
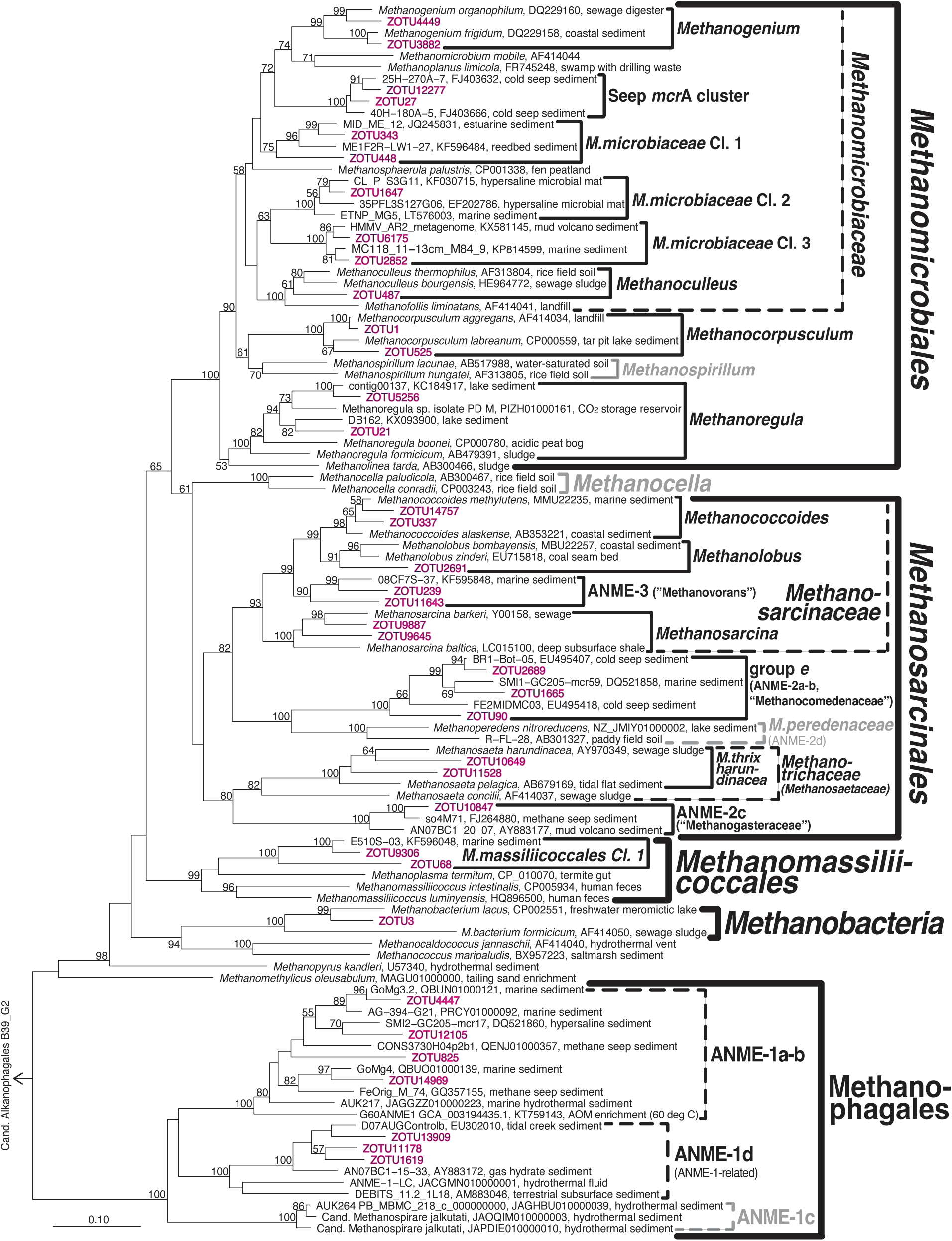
Phylogenetic tree of *mcr*A clades. Representative 97%-clustered ZOTUs of environmentally important clades based on read percentages are shown in magenta. Latin names of candidate taxa that have been proposed based on genomic analyses but lack pure culture isolates are placed in quotes. Clades that were not detected but included for illustration purposes are shown in gray. Note: ‘Hydrothermal ANME-1 cluster’ is treated as separate from ANME-1a-b in Figure 3 and the main text.

At the genus-and family-level we observe additional site-and geochemical zone-related trends (for phylogenetic terminology, see **Figure 4** and next section). AU1 is dominated by a new cluster of *Methanomicrobiales* (*Methanomicrobiaceae* Cluster 2) in surface sediments. Throughout the remaining core, sequences belonging to *Methanococcoides* and *Methanosarcina* (both *Methanosarcinaceae*) dominate, with significant contributions of ANME-1a-b, *Methanoregula* and *Methanomicrobiaceae* Cl. 1 in a few layers.

*Methanococcoides* and *Methanolobus* dominate sulfate reducing sediment of AU2 along with *Methanoregula* and *Methanogenium*. A major shift occurs in the SMT, where sequences of the “family-level” group *e* (Candidatus Methanocomedenaceae), a sister clade of anaerobic methane-oxidizing *Methanoperedenaceae* (both *Methanosarcinales*), dominate, along with methanotrophic ANME-2c archaea (one sample only). A second shift occurs below in the methanogenesis zone, toward a heterogeneous assemblage dominated by the new *Methanomicrobiaceae* Cluster 3, seep *mcr*A cluster (Lever & Teske, 2015), and *Methanoregula* (all *Methanomicrobiales*). Notably, *Methanothrix* (also known as *Methanosaeta*) account for significant percentages (∼10-15%) in several layers, as do sequences belonging to ANME-1d, a sister clade of ANME-1a-b (Lever et al., 2023); for further information, see next section). ANME-1a-b are also abundant at AU2, but mainly in and above the SMT.

*Methanococcoides*, *Methanosarcina*, *Methanogenium* and *Methanomicrobiaceae* Cl. 1, groups that were also abundant in the same zones of AU1 or AU2, dominate the bioturbation and upper sulfate reduction zone of AU3, along with *Methanocorpusculum*. As at AU2, there is a clear community shift in the lower sulfate reduction zone and SMT, where *mcr*A group *e*, ANME-2c, ANME-3 (“Candidatus Methanovorans”), seep *mcr*A cluster, and *Methanomicrobiaceae* Cl. 3 become dominant. In the lower part of the SMT the community shifts yet again, becoming dominated by ANME-1a-b, ANME-1d, and *Methanomassiliicoccales* in the methanogenesis zone.

The community zonation at AU4, which has no bioturbation zone, is distinct from the other sites. ANME-1-a-b dominates the sulfate-reduction zone and SMT and is replaced by ANME-1d as the dominant group in the upper part of the methanogenesis zone. The sulfate reduction zone and SMT additionally harbor significant percentages of *mcr*A group *e*, ANME-2c, and *Methanococcoides*, with *Methanogenium* and *Methanocorpusculum* being additionally present in surface sediment. In addition to ANME-1d, *Methanomassiliicoccales* Cluster 1 and to a lesser degree *Methanomicrobiaceae* Cl. 3 and ANME-2c account for significant percentages in the methanogenesis zone.

### mcrA phylogeny

An *mcr*A phylogenetic tree confirms the high diversity of methane-cycling archaeal taxa at the four sites (**Figure 4**). Most of this diversity is found in the *Methanomicrobiales* and *Methanosarcinales*. While metabolically well-characterized groups dominate the *Methanosarcinales*, uncharacterized environmental clusters within the *Methanomicrobiaceae* dominate the *Methanomicrobiales*. Phylogenetic diversity is considerably lower in the Candidate order Methanophagales, the *Methanomassiliicoccales* (class *Thermoplasmata*), and the *Methanobacteria*.

As already mentioned, the diversity of *Methanomicrobiales* is dominated by *Methanomicrobiaceae*, which include six of the eight *Methanomicrobiales* clusters present. Within the *Methanomicrobiaceae*, the genera *Methanogenium* and *Methanoculleus* have cultured members, whereas the Seep *mcr*A cluster and newly proposed *Methanomicrobiaceae* Clusters 1-3 are only known from environmental samples, including methane seeps (Seep *mcr*A cluster) and a range of marine sedimentary habitats (*Methanomicrobiaceae* Cl. 1-3; **Figure 4**). Since all cultured members of *Methanomicrobiaceae* perform methanogenesis by CO_2_ reduction using H_2_ or formate (and in some cases certain alcohols) as electron donors (Whitman et al., 2014), the four uncharacterized *Methanomicrobiaceae* groups may also perform these reactions. The remaining groups of *Methanomicrobiales* are represented by close relatives of *Methanocorpusculum aggregans* and a subcluster of *Methanoregula*. Cultured *Methanocorpusculaceae* reduce CO_2_ using H_2_ and formate, and in a few cases secondary alcohols, as electron sources (Whitman et al., 2014), while cultured *Methanoregula* reduce CO_2_ with H_2_, but not formate (exception: *Methanoregula formicica*; Yashiro et al., 2011).

The *Methanosarcinales* groups present are metabolically diverse. The methylotrophic *Methanococcoides* and *Methanolobus* grow by disproportionation of methanol and methylamines (both genera), and several additional C1 compounds (only certain *Methanolobus*; (Liang et al., 2022; Oremland & Boone, 1994). The closely related ANME-3 group is considered to be anaerobic methanotrophic (Bhattarai et al., 2019). *Methanosarcina* are substrate generalists, known to produce methane from H_2_/CO_2_, acetate, methanol, and methylamines, but not formate (Whitman et al., 2014). In addition, members of this group were more recently shown to grow by CO_2_ reduction via direct electron transfer (DIET) from minerals or syntrophic partners (Gao & Lu, 2021b; Rotaru et al., 2014). Outside of the *Methanosarcinaceae* the three additional groups of *Methanosarcinales* include *mcr*A group *e* (also known as ANME-2a). This group clusters at a high confidence-level with *Methanoperedenaceae*, members of which use nitrate-, iron(III)-, and manganese(IV) as electron acceptors for AOM (Ettwig et al., 2016; Haroon et al., 2013). Members of *mcr*A group *e* have been widely reported from methane seep environments with AOM (e.g., Hallam et al., 2003; Lloyd et al., 2006). In addition, the known methanotrophic ANME-2 group (also known as ANME-2c or group c-d; Knittel & Boetius, 2009) is present along with aceticlastic *Methanotrichaceae* (also known as *Methanosaetaceae*; Oren *et al*., 2014). Within the latter, all sequences fall into a genus-level cluster with the previously isolated *Methanothrix pelagica* and *Methanothrix harundinaceae*. Notably, *Methanothrix harundinacea*, similar to certain *Methanosarcina*, were shown to also grow by CO_2_ reduction via DIET (Gao & Lu, 2021a).

The community structure of *Methanophagales* consists of two groups. The ANME-1a-b, which is one of the most studied groups of methanotrophic archaea. Different from phylogenomic or 16S rRNA gene sequence analyses, ANME-1a cannot be reliably separated from ANME-1b based on *mcr*A sequence analyses (hence the name ANME-1a-b). In addition to ANME-1a-b, we detect another major branch of *Methanophagales*, belonging to the recently named ANME-1d cluster (Lever et al., 2023). This cluster, which is also known as ANME-1-related group (Aromokeye et al., 2020; Lever & Teske, 2015), has been found across subseafloor sediments (Lever, 2008), hydrothermal vents with serpentinitic fluids (Kelley et al., 2005), deep subsurface coalbeds (Fry et al., 2009), and gas hydrate sediments (Kormas et al., 2008). Frequently, the ANME-1d have been assumed to be anaerobic methanotrophs same as ANME-1a and ANME-1b. Yet, ANME-1d is phylogenetically clearly distinct from ANME-1a-b (**Figure 4**), and has even been proposed to constitute a separate order based on *mcrA* sequence (dis)similarity to ANME-1a-b, Lever & Teske, 2015; also see Discussion).

The *Methanomassiliicoccales* detected at the four sites all belong to an uncharacterized phylogenetic cluster that is distinct from the cultured genus *Methanomassiliicoccus*, or the genome-sequenced candidate genera *Methanoplasma* or *Methanomethylophilus*. The closest relatives of this cluster based on mcrA phylogeny were detected in other (marine) sedimentary environments (e.g., KF596048, KF595850, AND KF595354 from Zhou et al., 2015). Pure culture and genomic evidence suggest these three genera to produce methane by reducing methanol or methylamines with H_2_ (Kröninger et al., 2017; Lang et al., 2015). Members of the CO_2_-reducing *Methanobacteriales*, which were only detected at significant abundances in one sample from the bioturbation zone of AU2, are represented by one phylotype (ZOTU3) that is most closely related to *Methanobacterium lacus*.

### Substrate use

The substrate use by different groups of methane-cycling microorganisms provides further insights into their metabolic/catabolic pathways in sediment (**Figure 5**). Taxa that use acetate or methylated compounds without H_2_ were largely restricted to the bioturbated and sulfate-reducing zones of AU1-AU3. Taxa that use methanol+H_2_, on the other hand, were largely absent from bioturbated and sulfate-reducing sediments, instead increasing to significant percentages in methanogenic sediments. Known CO_2_-reducing taxa showed rather patchy distributions, dominating AU2 except in the SMT, showing locally high percentages in all major zones of AU3, and being numerically abundant in surface and a few deeper samples of AU1 and AU4. Putative anaerobic methanotrophs (ANME-1a-b, Hydrothermal ANME-1, ANME-2) accounted for high percentages in all SMTs, but only dominated the SMT of AU4. In addition, methanotrophs dominated the methanogenesis zone of AU3 and sulfate-reduction zone of AU4, and were present at locally significant contributions in sulfate-reducing and bioturbated layers of AU1-AU3. Remarkably, the energy substrates of a major, locally even dominant, fractions of *mcr*A reads are unknown. These dominant, physiologically uncharacterized groups are ANME-1 in methanogenic sediments of AU2 to AU4 and *mcr*A group *e* in the SMT of AU2.

**Figure 5.**
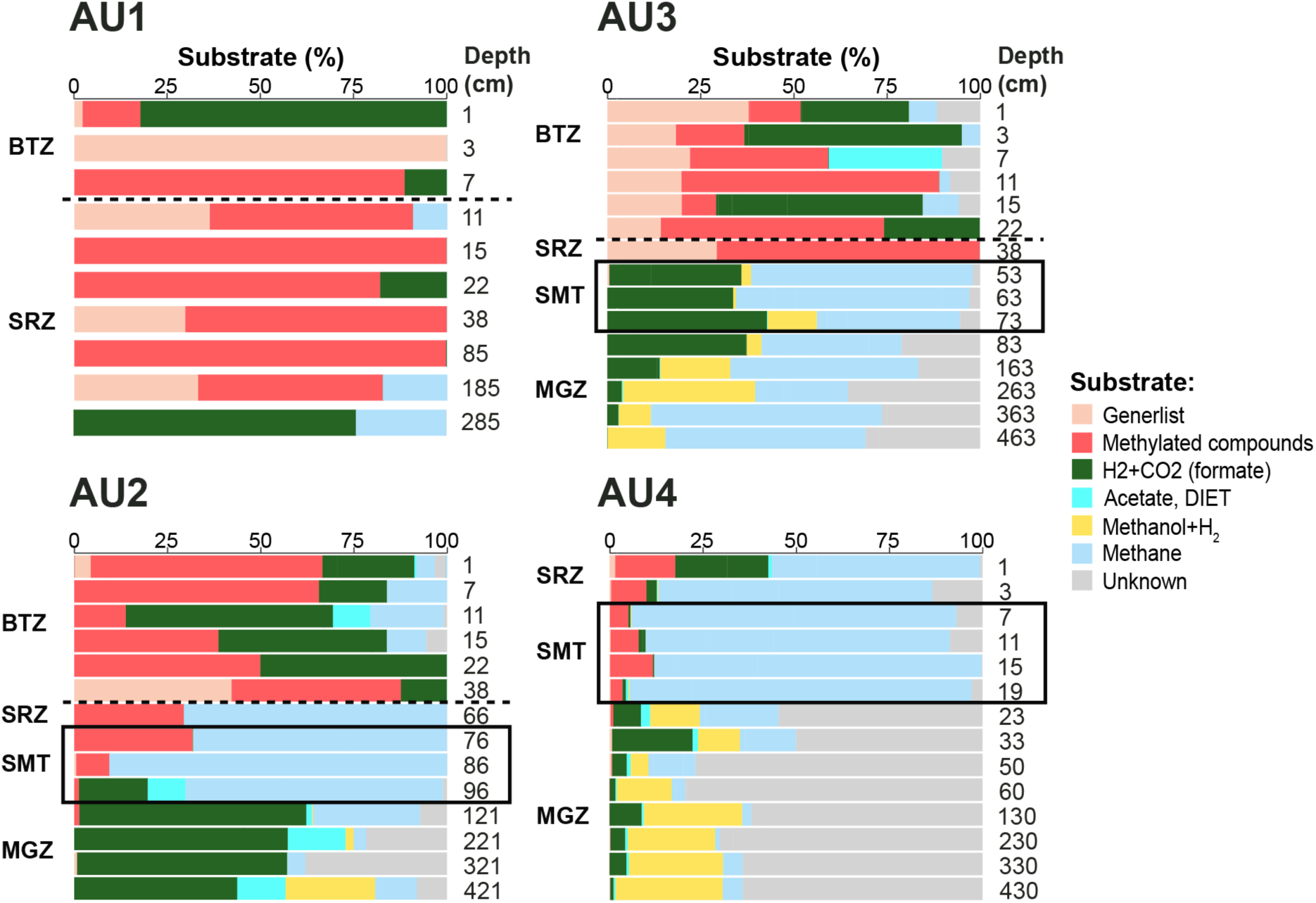
Depth profiles of major energy substrates at AU1-AU4 inferred from taxonomic identity, and related physiological knowledge, of major *mcr*A clades. Samples above the horizontal dashed lines were located within the bioturbation zone. Black boxes indicate samples that were located within the sulfate-methane transition. [Abbreviations: BTZ = bioturbation zone, SRZ = sulfate-reducing zone, SMT = sulfate-methane transition, MGZ = methanogenic zone].

### Zonation of mcrA community structure at the ZOTU-level

Despite the site-specific differences in dominant clades, methane-cycling archaea at the ZOTU-level show clear zonation. ZOTU-level community fingerprints show a clear relationship with Sulfate concentrations (NMDS1; **Figure 6A**). In addition, site-specific clustering can be observed (NMDS2), with strong overlaps in methane-cycling archaeal communities between AU1, AU2, and AU3, but more distinct communities in the sulfidic, more organic carbon-rich sediments of AU4. Similar patterns can be observed in relation to biogeochemical zone (**Figure 6B**). Communities in sediments from the bioturbation zone overlap with those in SRZs. Communities in SMTs show considerable variation, depending on location clustering with samples from the BTZ, SRZ, or MGZ. By contrast, samples from the methanogenesis zone are clearly distinguished from bioturbated or sulfate-reducing sediment at the ZOTU-level.

**Figure 6.**
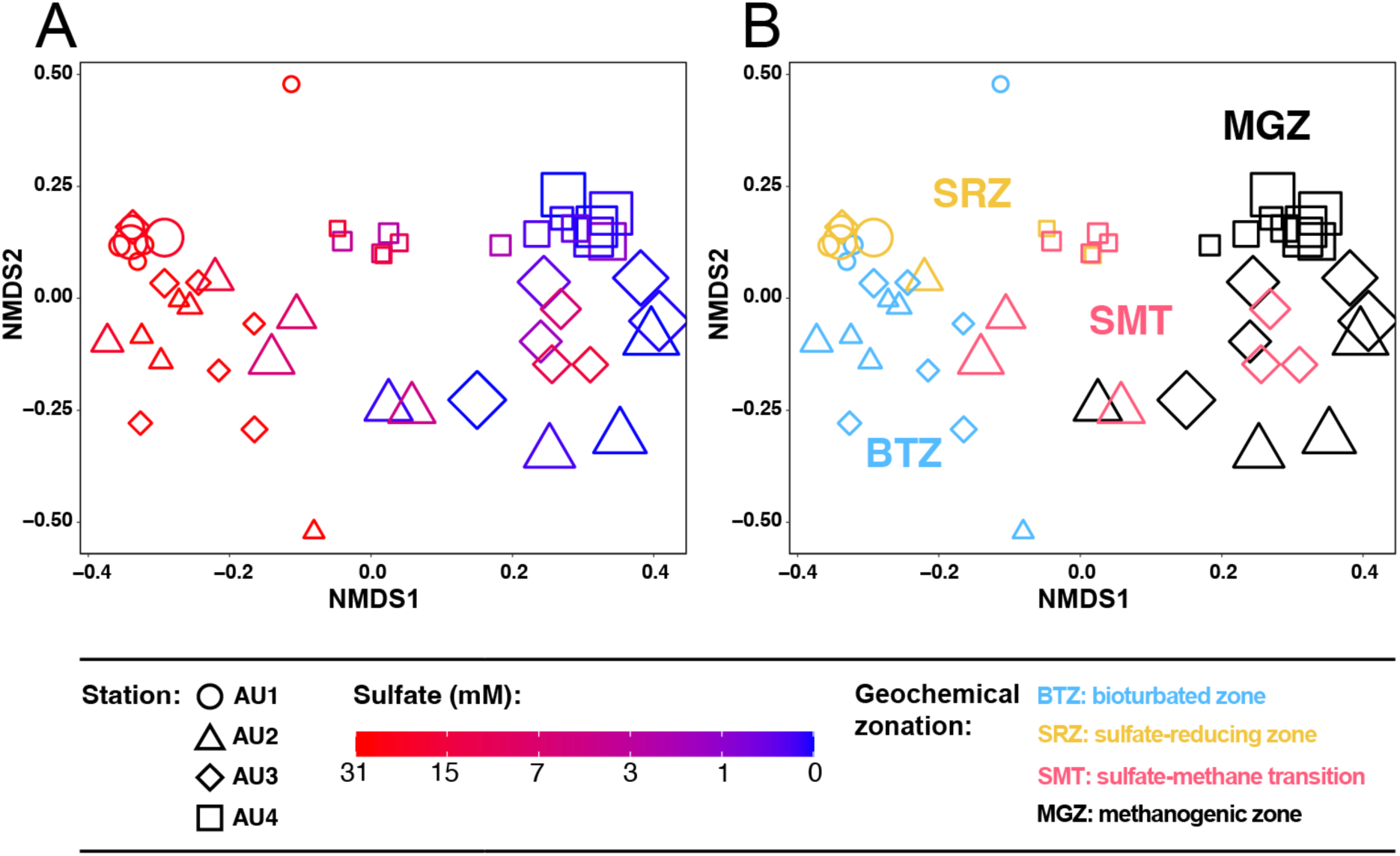
Non-Metric Multidimensional Scaling (NMDS) based on Bray-Curtis distances of *mcr*A communities at the ZOTU-level in relation to (**A**) sulfate concentrations and (**B**) biogeochemical zone.

## Discussion

Marine sediments are globally important reservoirs of methane, with the bulk of this methane being internally produced and consumed by methane-cycling archaea. While a considerable body of literature exists on advective systems, little is known about the environmental drivers of methane-cycling archaeal community structure and metabolism in diffusion-dominated sediments, where most marine methane production and consumption takes place. By integrating geochemical, stable isotopic, and methane-cycling archaeal abundance and community data from coastal eutrophic to off-shore oligotrophic sites, we explore the drivers of methanogenesis and AOM in continental margin sediments. Our analyses indicate that active methane-cycling is not restricted to sulfate-methane transitions (SMTs) and methanogenesis zones (MGZs), but also occurs widely in bioturbation zones (BTZs) and sulfate-reduction zones (SRZs). Pervasive vertical and horizontal changes in methane-cycling communities and inferred metabolisms indicate a major role of depth-and site-specific environmental variables in driving the methane cycle in continental margin sediments.

### Distribution of methanogenic and methanotrophic activity

Downcore concentration profiles of methane, with low (micromolar) concentrations throughout the BTZ and SRZ, increasing to millimolar concentrations where sulfate is depleted in deeper layers, are consistent with the standard biogeochemical zonation (e.g., Jørgensen & Kasten, 2006) (**Figure 2**). Accordingly, methanogenesis is suppressed by microbial respiration with higher energy yields, e.g. by sulfate reduction and by metal reduction, in surface sediments, where sulfate and metal oxides are present. The distinct increase in methane concentrations in deeper, sulfate-depleted layers suggests that methanogenesis only becomes the dominant respiration reaction once energetically superior electron acceptors have become largely consumed. The almost complete absence of methane in the SMT is, moreover, consistent with sulfate-dependent AOM consuming upward-diffusing methane from the MGZ in sediment layers where this methane meets downward-diffusing sulfate from seawater.

While the concentration profiles of sulfate and methane are consistent with peak methane-cycling activity in the SMT and MGZ, isotopic data suggest that active methane cycling, with *in situ* methane oxidation and possibly production, is also present in the BTZ and SRZ. At AU1 and AU2, the δ^13^C-CH_4_ becomes increasingly more positive from the MGZ into the overlying SRZ and BTZ. Gradual oxidation of methane that has escaped oxidation in the SMT and is diffusing upward into the SRZ may drive this shift to higher δ^13^C-CH_4_. Yet, the limited amount of isotopically “light” methane that is oxidized to DIC is too small to significantly lower the δ^13^C-DIC, given the much higher input of DIC from organic matter mineralization.

In addition, we cannot rule out that methanogenesis is co-occurring with AOM throughout the SRZ and BTZ. The nearly parallel δ^13^C-CH_4_ and δ^13^C-DIC profiles throughout AU1, and in certain surface sedimentary intervals of AU2 and AU3, are consistent with low rates of CO_2_ reduction. The δ^13^C-CH_4_ may follow the δ^13^C-DIC due to constant isotopic fractionation during the conversion of CO_2_ (DIC) to methane. Similarly, low rates of other methanogenic reactions, e.g. acetate disproportionation and methyl group dismutation, cannot be ruled out based on the C-isotopic data given the absence of isotopic data on acetate and methylated compounds and the fact that fractionations produced by these reactions could be masked by other C-isotopic fractionations related to methane-cycling.

Further down, the dominant processes involved in methane cycling become more evident. In the SMTs of AU2 and AU3, the depletion of sulfate and increase in methane, which coincide with slight increases in δ^13^C-CH_4_ and strong decreases in δ^13^C-DIC, is a typical sign of AOM. At both sites δ^13^C-DIC values drop below those of δ^13^C-TOC, indicating oxidation of isotopically light methane as a major source of DIC in these layers. Notably, AU4 shows a different trend. Despite the strong upward decrease in methane concentrations across the SMT, the δ^13^C-CH_4_ and δ^13^C-DIC both show upward, parallel decreases. We propose that at AU4 significant rates of methanogenic CO_2_ reduction co-occur with AOM within the SMT. Despite the clear decrease in methane concentrations from the MGZ through the SMT, which suggests net oxidation of methane, the stronger negative isotopic fractionation associated with methanogenic CO_2_ reduction appears to dominate δ^13^C-CH_4_ values over the comparatively weaker isotopic fractionation associated with AOM. This interpretation matches environmental measurements of C-isotopic fractionations, which indicate considerably higher C fractionations associated with methanogenic CO_2_ reduction (α=1.045-1.082; reviewed in Conrad, 2005) than with AOM (α=1.004-1.021; reviewed in Alperin & Hoehler, 2009). A similar co-occurrence of methane production and AOM in the SMT was previously proposed for organic-rich coastal sediments based on radiotracer incubations (Beulig et al., 2019) and C-isotopic analyses of methane-cycling microbial aggregates within SMTs (Alperin & Hoehler, 2009). In addition, given the shallow sediment depth of the SMT at AU4 (∼10 to 20 cm), it is possible that the strong decrease in methane concentration near the sediment surface is not solely caused by AOM, but also by ebullition of methane gas bubbles into overlying water.

Measured δ^13^C-CH_4_ and δ^13^C-DIC profiles in the MGZs of AU2 through AU4 are nearly parallel and increase with depth, consistent with CO_2_ reduction as the dominant methanogenic pathway. The offset between δ^13^C-CH_4_ and δ^13^C-DIC is remarkably constant between sites, mostly ranging from -68.2 to -80.5 per mil, which corresponds to an isotopic fractionation factor (α) of 1.075-1.085 for methanogenic CO_2_ reduction. These values are among the highest reported from marine sediments (Conrad, 2005). At all three sites, the rates of methanogenic CO_2_ reduction are sufficiently high to drive δ^13^C-DIC into the positive range, to values that are up to -40 per mil lower than the δ^13^C-TOC (AU4).

### Thriving of methane-cycling archaea in response to vertical geochemical gradients

Copy numbers of *mcr*A are similar in surface sediments of all sites but are uniformly very low in absolute abundances (10^2^ to 10^4^ copies g^-1^ sediment) and relative abundances (mcrA:16S rRNA gene ratios of 10^-5^-10^-4^ (∼0.001-0.01%) (**Figure 3A**). Comparing deeper layers, there is, however, a clear increase in copy numbers from the oligotrophic AU1 to the eutrophic AU4 (**Figure 3A**). This trend in deeper layers matches the increase in sedimentary organic carbon content and reactivity from the deep Norway Trench site (AU1, 586 m) to the southern slope of the Norway Trench (AU2, 319 m) and shallow shelf sites, which include the sandy, metal-reduction dominated Kattegat site (AU3, 43m) and muddy, organic-rich, sulfidic Lillebælt site (AU4, 37 m; Kristensen et al., 2018; Deng et al. 2020). Presumably an increase in energy availability due to higher organic matter content and reactivity drives this increase in subsurface methane-cycling archaeal abundances from offshore to nearshore. Nevertheless, ratios of *mcr*A to total 16S rRNA gene copy numbers indicate that methane-cycling archaea only account for a small fraction (mostly <<1%) of the total community, even in methanogenic sediments of AU4 (**Figure 3A**). Thus, methane-cycling archaea appear to be part of a rare, albeit geochemically important, microbial biosphere in continental margin sediments.

In addition to the trends in *mcr*A copy numbers across sites, there are clear vertical trends within sites. At AU2 through AU4, *mcr*A copy numbers increase by an order of magnitude from the lower SRZ to the SMT and uppermost layer of the MGZ, indicating thriving of methane-cycling archaeal populations in response to more favorable geochemical regimes (**Figure 3A**). Thus, methane-cycling archaea appear to be an exception to the notion that microbial populations in subsurface sediments below the bioturbation zone generally are too energy-limited to thrive, consisting of surviving, at most self-maintaining populations (Bradley et al., 2020). This interpretation is in line with research on deep subseafloor sediments of the Peru Trench, where *mcr*A was below detection in sulfate reducing sediment, but became widely detectable in deeper layers of the sulfate-depleted SMT and MGZ (Lomstein et al., 2012; Lever et al., 2023).

### Methane-cycling archaeal communities of bioturbated and sulfate-rich sediments

Phylogenetic analyses reveal a diverse methane-cycling community that varies significantly with location and in relation to vertical biogeochemical zones. Herein the biggest changes, both at the ZOTU-level and at higher phylogenetic levels, occur between sulfate-rich (surface) sediments (BTZ, SRZ), the SMT, and the MGZ (**Figure 3B-C**; **Figure 6**). These community changes are not only apparent at the relative abundance-level, but also when taxon-specific absolute abundances are compared across vertical biogeochemical zones (**Figure 7**).

**Figure 7.**
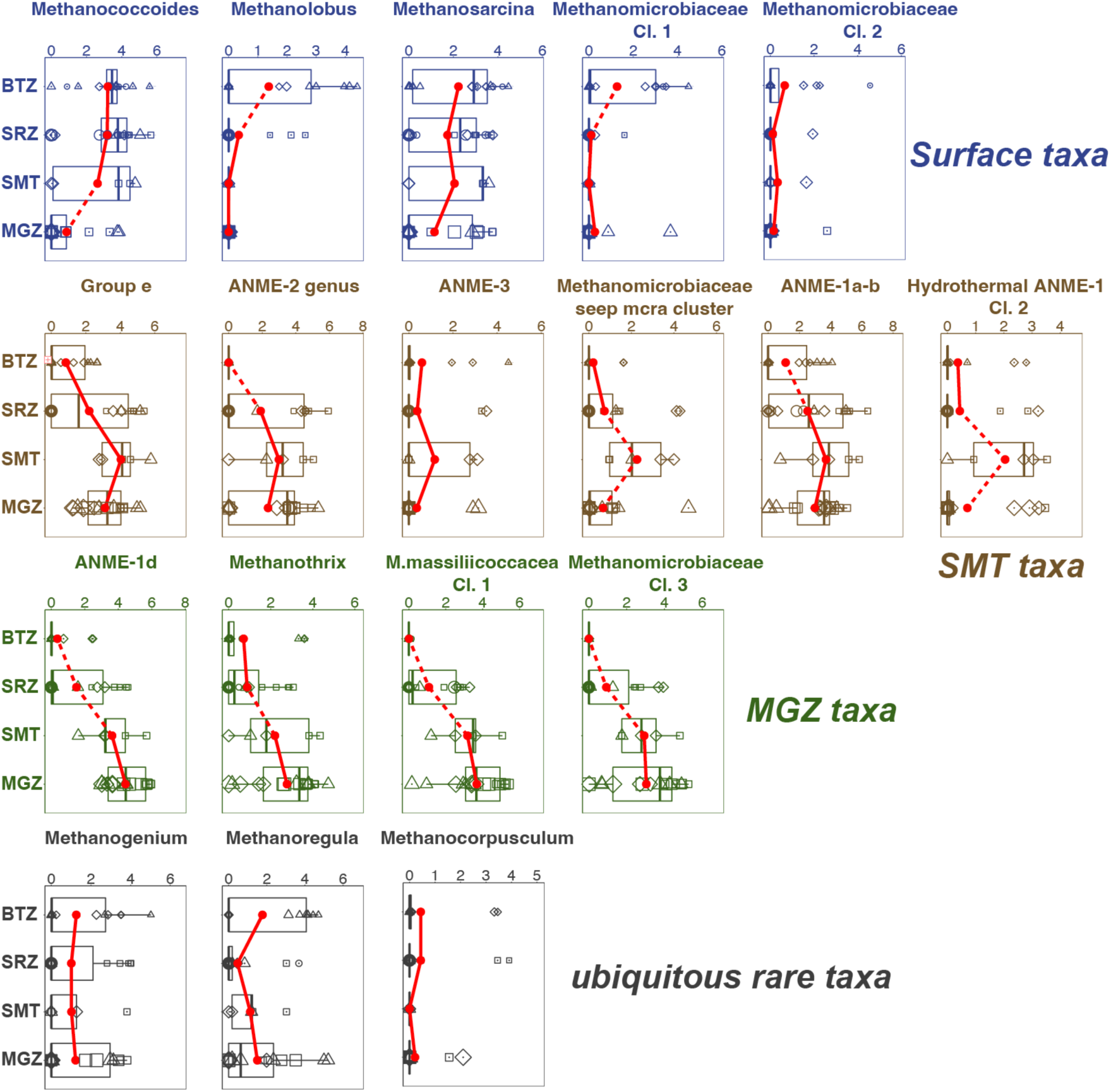
Vertical trends in the absolute abundances of different methane-cycling archaeal groups in relation to geochemical zone (BTZ=bioturbation zone; SRZ=sulfate reduction zone; SMT=sulfate-methane transition; MZ=methanogenesis zone). Absolute abundances of *mcr*A genes (log_10_ copies g^-1^ dry weight) for each group in each sample were calculated by multiplying total *mcr*A copy numbers by the percent contribution to mcrA reads by that group. We observe four distinct trends, which we classified into four categories. *Surface taxa, SMT taxa*, and *MGZ taxa* refer to methane-cycling archaea with highest abundances in bioturbation or sulfate reduction zones (*surface taxa*), sulfate-methane transitions (*SMT taxa*), and methanogenesis zones (*MGZ taxa*). A fourth category (*ubiquitous rare taxa*) showed no clear depth trend and was characterized by low abundances. The mean values of *mcr*A genes within each geochemical zone are indicated by the red dots, boxes indicate a 50% confidence interval. Dashed lines connecting adjacent points indicate significant differences of mean values (Wilcoxon test). [Note: Copy numbers are expressed in per gram dry weight to eliminate effects of sediment compaction on *mcr*A copy numbers.]

One of the most striking phylogenetic trends across the study sites is the shift from high percentages of *Methanosarcinales* in BTZ, SRZ, and SMT sediments to much lower contributions in MGZs. This indicates that higher sulfate concentrations lead to higher contributions of *Methanosarcinales*. The reasons are offered by the vertical phylogenetic (and metabolic) zonation of individual Methanosarcinales groups. Specialized, methyl-dismutating *Methanococcoides* and *Methanolobus* and generalistic *Methanosarcina* dominate both the BTZ and SRZ and decrease in both relative and absolute abundances with depth (**Figures 3C and 7**). The fact that these taxa, which share the metabolic potential for methyl-dismutation, dominate methanogenic communities in sulfate-rich sediments matches the notion that methylated compounds, such as methanol, methyl sulfides, and methylamines, are not utilized by most competing respiratory organisms. This enables these methylotrophic methanogens to thrive in sediments where sulfate or metal reduction dominate respiration (Maltby et al., 2016; Xiao et al., 2018; Zhuang et al., 2018).

In addition to *Methanosarcinales*, *Methanomicrobiales* account for a significant, in some places dominant, fraction of the methane-cycling archaeal community in sulfate-rich layers, with the highest percentages found in the bioturbation zone (**Figure 3C**). All cultivated members of this order, which include the genera *Methanogenium*, *Methanocorpusculum*, and *Methanoregula* in the BTZs of AU2 through AU4, are obligate CO_2_-reducing methanogens (Whitman et al., 2014; Garcia et al., 2017). Presumably, the novel genus-level lineages *Methanomicrobiaceae* Clusters 1 and 2, which were mostly found from AU1 to AU3, and which had their highest absolute abundances in BTZs (**Figure 7**), are no exception. The significant percentages of *Methanomicrobiales*, and of *Methanosarcina* which also include facultative CO_2_ reducers, are in line with δ^13^C-CH_4_ and δ^13^C-DIC trends that suggest CO_2_ reduction in the BTZ and SRZ, but goes against the notion that CO_2_-reducing methanogens are outcompeted by sulfate and metal reducers for H_2_ in these sediments (Hoehler et al., 1998; Lovley & Goodwin, 1988). A potential explanation is redox oscillations caused by macrofaunal ventilation and seawater advection, or episodic sediment mixing and import of labile organic matter by macrofaunal reworking. These fluctuations in geochemical conditions may prevent sulfate and metal reducers from drawing H_2_ concentrations down to steady-state levels that are too low to energetically support methanogenic CO_2_ reduction. Alternatively, it is possible that CO_2_ reducers do not rely on H_2_, but instead use electrons from DIET, supplied by syntrophic partner organisms, to reduce CO_2_. CO_2_ reduction via DIET has been shown in laboratory experiments with *Methanosarcina* (Holmes et al., 2018) and was recently proposed to dominate CO_2_ reduction by *Methanomicrobiales* in lake sediments (Meier et al., 2024).

Despite the decrease in methane concentrations from the SMT to the seafloor (**Figure 2**) and isotopic and genetic evidence for an active methane cycle in the BTZ and/or SRZ, only small subpopulations of putatively methanotrophic archaea appear to be supported (0 to 20% of *mcr*A reads) at AU1-AU3 (mainly ANME-1a-b; **Figure 3**). Potential reasons include the low net energy gains from sulfate-dependent AOM, which in many cases involves energy partitioning between methanotrophic archaea and sulfate-reducing partner organisms. In addition, it cannot be ruled out that, in the BTZs of AU1 through AU3, which receive episodic input of O_2_ and nitrate due to macrofaunal ventilation (Deng et al., 2020), physiologically more resilient methane-oxidizing bacteria (MOB) successfully compete with ANMEs for methane. This possibility, which has also been proposed for lake sediments (van Grinsven et al., 2022) is underscored by clear outnumbering of ANMEs by MOBs based on functional gene copy numbers. Copy numbers of the alpha subunit of partial methane monooxygenase (*pmo*A), a key gene of aerobic methane oxidation, are in the range of 10^5^ to 10^7^ copies g^-1^ sediment in the BTZs, and thus 10^4^-to 10^6^-fold higher than *mcr*A copy numbers of ANMEs (**Figure 7**). An exception to the notion is the non-bioturbated, mostly sulfidic surface sediment of AU4. Here ANME-1a-b and ANME-2 collectively account for >50% of *mcr*A reads in the SRZ. We propose that due to the high methane flux and shallow depth of the SMT (5 to 20 cm) at this site, significant amounts of methane escape oxidation in the SMT and are, at least partially, consumed by ANMEs in the overlying only 5 cm thick SRZ.

### Methane-cycling archaeal communities of sulfate-methane transitions

We detect all known ANMEs, except *Methanoperedenaceae*, in the sediments studied. While there are strong taxonomic overlaps between sites, each of the three SMTs is dominated by a different ANME group. AU2 is dominated by ANME-2a-b (Candidatus Methanocomedenaceae), AU3 by ANME-2c (Candidatus Methanogasteraceae; both Methanosarcinales), and AU4 by the ANME-1a-b family (Candidatus Methanophagaceae; Candidatus Methanophagales). Moreover, while ≥80% of *mcr*A read percentages at AU2 and AU4 belong to ANMEs, only about half of the reads in the SMT of AU3 belong to ANMEs. The other half consist largely of uncultured *Methanomicrobiales* (seep *mcr*A cluster, *Methanomicrobiaceae* Cl. 3).

The dominance of anaerobic methanotrophs in SMTs is expected, and the preference of these groups for SMTs is supported by the fact that absolute abundances of all ANME groups except ANME-1d (*discussed in next section*) were highest in SMTs (**Figure 7**). Yet, the reasons for the distinct differences in dominant groups between locations are not clear. The fact that ANME-2a-b, ANME-2c, and ANME-1a-b co-occur in significant percentages in each location (**Figure 3**) argues against competitive exclusion over a limiting resource, and suggest that instead niche differences combined with location-specific environmental variables may cause dominance of different groups. While previous studies have suggested that ANME-2 thrive at higher sulfate concentrations than ANME-1 (Knittel & Boetius, 2009; Yanagawa et al., 2011), we observe significant contributions of ANME-1a-b also in the SRZ of AU4 and within the BTZ of AU2 (**Figure 2**). Given that iron and manganese reduction (AU2) and iron reduction (AU3) are dominant respiratory reactions at two of our sites (Kristensen et al., 2018), AOM coupled to metal reduction may also take place and locally select for ANME-2a-b and ANME-2c taxa with the capacity for this form of AOM. While ANME-1 and ANME-2 strongly overlap in distributions within SMTs, and even in SRZs, only ANME-1a-b and its sister clades ANME-1d and Hydrothermal ANME-1 Cluster 2 were detected in high abundances throughout deeper, sulfate-depleted MGZs of AU3 and AU4. This matches the reduced dependence of ANME-1 on sulfate as an electron acceptor (Yanagawa et al., 2011) and the notion that certain ANME-1 are (facultative) methanogens (Beulig et al., 2019; Lever et al., 2023; Lloyd et al., 2010).

Even though the sulfate-methane concentration gradients indicate that almost all methane produced in the MGZs of AU2-AU4 is consumed in the overlying SMTs, there are several inconsistencies between our isotopic and genetic data. For instance, while AU3 has C-isotopic profiles indicating that AOM is by far the dominant methane-cycling process, we detect relatively high abundances of putatively CO_2_-reducing *Methanomicrobiales*. An explanation could be that cell-specific activities of ANMEs are far higher than those of *Methanomicrobiales*, or that the latter are dormant. Yet, at least the notion of dormancy contradicts the absolute abundances of the uncultured Seep *mcr*A cluster, which is the dominant Methanomicrobiales group in SMTs and indicates a clear preference and even growth stimulation of this group within SMTs (**Figure 7**). Given that the Seep *mcr*A cluster is often detected in hydrocarbon and methane seeps with AOM (Lazar et al., 2012; Lever & Teske, 2015), the possibility of this group engaging in AOM should be investigated.

On the opposite side of the spectrum, the isotopic data at AU4, with parallel gradients in δ^13^C-CH_4_ and δ^13^C-DIC across the SMT, indicate that CO_2_ reduction is taking place in parallel to AOM, and even overriding the isotopic imprint of AOM. This CO_2_ reduction could be performed, at least in part, by ANME-1a-b, consistent with previous radiotracer-based evidence for CO_2_ reduction by this group in SMTs (Beulig et al., 2019). In addition, or alternatively, the less studied, ANME-1d group, which is also abundant in the SMT of AU4, could be involved in CO_2_ reduction. This would match the generally deeper distribution of this group compared to ANME-1a-b (Figure 3C), and the fact that this group dominates methanogenic subsurface sediments in other locations (Lever et al., 2023).

### Methane-cycling archaeal communities of methanogenesis zones

While δ^13^C-CH_4_ and δ^13^C-DIC profiles suggest CO_2_ reduction as the dominant methanogenic pathway in the MGZs of AU2 through AU4, taxonomic compositions paint a confusing picture (**Figure 3C**). Known CO_2_-reducing methanogens (*Methanoregula*) dominate in only a single sample from AU2, while all other samples are dominated by uncultured Methanomicrobiales, putative methane oxidizers of the ANME-1 group (ANME-1a-b, ANME-1d, Hydrothermal ANME-1 Cl. 2) and a new family-level cluster of *Methanomassiliicoccales*. When absolute abundances are considered, then at least ANME-1d, the *Methanomassiliicoccales* cluster, *Methanomicrobiaceae* Cl. 3, and the less abundant aceticlastic genus *Methanothrix* are likely methanogens (**Figure 7**). *mcr*A copy numbers of these groups increase ∼100 to 10,000-fold from BTZ and SRZ sediments to more methane-rich layers of the SMT and MGZ, and additionally are highly correlated with methane concentrations (**Figure S1**; all with Spearman’s Rho>0.6, p<0.001,).

The fact that methanogenic sediments at all three sites are dominated by uncultured groups highlights the need for more cultivation research on marine methanogens. Even within the *Methanomicrobiales*, which include 24 published isolated species, only 5 of these isolates (*Methanogenium cariaci*, *M.genium marisnigri*, *M. organophilum*, *Methanolacinia paynteri*, *Methanoculleus thermophilicum*) are from marine environments (Garcia et al., 2017). The fact that the 3 dominant *Methanomicrobiales* taxa in methanogenic sediments belong to uncultured groups only underscores the need for more cultivation efforts. Nevertheless, given that all 5 marine Methanomicrobiales isolates were CO_2_-reducers of the family *Methanomicrobiaceae*, which our three uncultured groups also belong to (**Figure 4**), it seems that our uncultured groups also include CO_2_-reducing methanogens.

The uncertainty increases when the dominant groups of ANME-1 are examined. While ANME-1a-b the *mcr*A copy number peak in the SMT is consistent with methanotrophy (**Figure 7**), the consistently high copy numbers and read contributions in MGZs support the notion that this group includes facultative CO_2_-reducing methanogens (Beulig et al., 2019; Lever et al., 2023; Lloyd et al., 2011). A similar case could be made for the Hydrothermal ANME-1 Cl. 2. This group, which was first classified in hydrothermal sediment of Guaymas Basin (Lever & Teske, 2015) – despite also having an *mcr*A copy number peak in the SMT -shows an increase in read contributions from the SMT to the MGZ of AU3. The conditions under which either group might switch to a methanogenic lifestyle are unclear but do not appear to be strictly coupled to biogeochemical zone, as shown by radiotracer experiments indicating ‘cryptic’ CO_2_ reduction by ANME-1a-b in SMTs with net methane oxidation (Beulig et al., 2019).

While the distributions of ANME-1a-b and Hydrothermal ANME-1 Cl. 2 support the idea of these groups being facultative methanogens, the ANME-1d group shows abundance distributions that indicate a primarily, if not solely, methanogenic lifestyle. This group not only increases in read percentages from SMTs to MGZs of AU2-AU4, and becomes the dominant group in the MGZ of AU4 (**Figure 3C**), but also has peak *mcr*A copy numbers in MGZs that are mostly simply explained by a methanogenic lifestyle (**Figure 7**). Any inferences regarding methanogenic pathway deserve caution in the absence of cultivation or genomic data on ANME-1d. Yet, C-isotopic signatures indicative of CO_2_ reduction as the dominant methanogenic pathway at AU4, where ANME-1d account for 60% of *mcr*A reads, are clearly in line with this group carrying out methanogenic CO_2_ reduction at the sites studied. Our interpretation matches results from deep subseafloor sediments in which ANME-1d dominated methane-cycling archaeal communities in layers that were >100m below the SMT and had been buried below the depth of sulfate-depletion for at least 400,000 years (Lever & Teske, 2015).

Another striking observation is the strong *mcr*A copy number increase of an unclassified *Methanomassiliicoccales* cluster in MGZs. This group was below detection in the majority of BTZ samples, increased slightly within the SRZ, and then increased by three orders of magnitude across the SMT into the MGZ (**Figure 7**). While this novel group remains physiologically uncharacterized, the phylogenetic clustering within the Methanomassillicoccales suggests that methanogenic methyl group reduction with H_2_ as a likely metabolism (Kröninger et al., 2017; Lang et al., 2015). Remarkably, the increase in absolute abundances of this putatively methanol-reducing group correlates negatively with the strong depth-related decrease in methyl-dismutating taxa (*Methanococcoides, Methanolobus, Methanosarcina*, *p*<0.05). This suggests a switch from methyl dismutation as the prime methylotrophic methanogenic pathway in sulfate-rich surface sediments to methyl-reduction in deeper, sulfate-depleted layers (for further analyses, see next section). With mcrA copy numbers in the MGZ that are only second to ANME-1d and mcrA read percentages exceeding 20% in many of the deeper methanogenic layers, we propose that methyl group reduction is an important, widely overlooked methanogenic pathway in deep methanogenic marine sediments.

### Drivers of methanogenic pathway distributions

The observed taxa distributions suggest environmental variations in the distributions of methanogenic and anaerobic oxidation of methane (**Figures 3 and 4**) that are not always visible within measured chemical concentration profiles and isotopic compositions (**Figure 2**) due to their cryptic recycling and lack of a clear geochemical imprint. To explore potential drivers behind these distributions, we next take a thermodynamic approach with a focus on *in situ* energy yields of various reactions.

Given that H_2_ concentrations were not determined as part of this project, we test two H_2_ concentration scenarios. These H_2_ concentrations are in the range of the lowest ones found in anoxic sediments (0.1 nM) and in the typical range of methanogenic sediment (10 nM) (Hoehler et al., 1998; Lovley & Goodwin, 1988). We calculate that CO_2_ reduction is never exergonic at H_2_ concentrations of 0.1 nM, but always exergonic at concentrations of 10 nM, with Gibbs energies in the range of -40 to -12 kJ mol^-1^ (**Figure 8A**). Herein the most negative (exergonic) values were calculated for BTZs and SRZ, while the least negative (lowest energy yielding) values (-20 to -10 kJ mol^-1^ reaction) were calculated for MGZs. Gibbs energy values for MGZs were thus close to the biological energy quantum (-10 kJ mol^-1^; (Hoehler et al., 2001) in a range previously determined for methanogens in MGZs (Hoehler, 2004), suggesting that H_2_ concentrations in the range of 10 nM H_2_ are plausible for the MGZs of all sites. That is assuming that CO_2_ reduction is indeed proceeding through H_2_ as the main electron carrier, and not via DIET as recently proposed for lake sediments (Meier et al., 2024). The scenario changes in surface and sulfate-rich sediments, where due to lower concentrations of the end product methane, Gibbs energies are more negative. We estimate that H_2_ concentrations around 1 nM, i.e. halfway between Gibbs energies for [H_2_]=0.1 nM and [H_2_]=10 nM in **Figure 8A**, are sufficient to support methanogenic CO_2_ reduction in these layers. Notably also, the Gibbs energies of CO_2_ reduction are in the same range in SMTs as in MGZs, suggesting that at ∼10 nM H_2_ methanogenic CO_2_ reduction is thermodynamically favorable in SMTs. This could explain the isotopic evidence for CO_2_ reduction in the SMT of AU4, and suggests that CO_2_ reduction by ANME-1a-b deserves consideration within the SMTs of AU2 and AU3.

**Figure 8.**
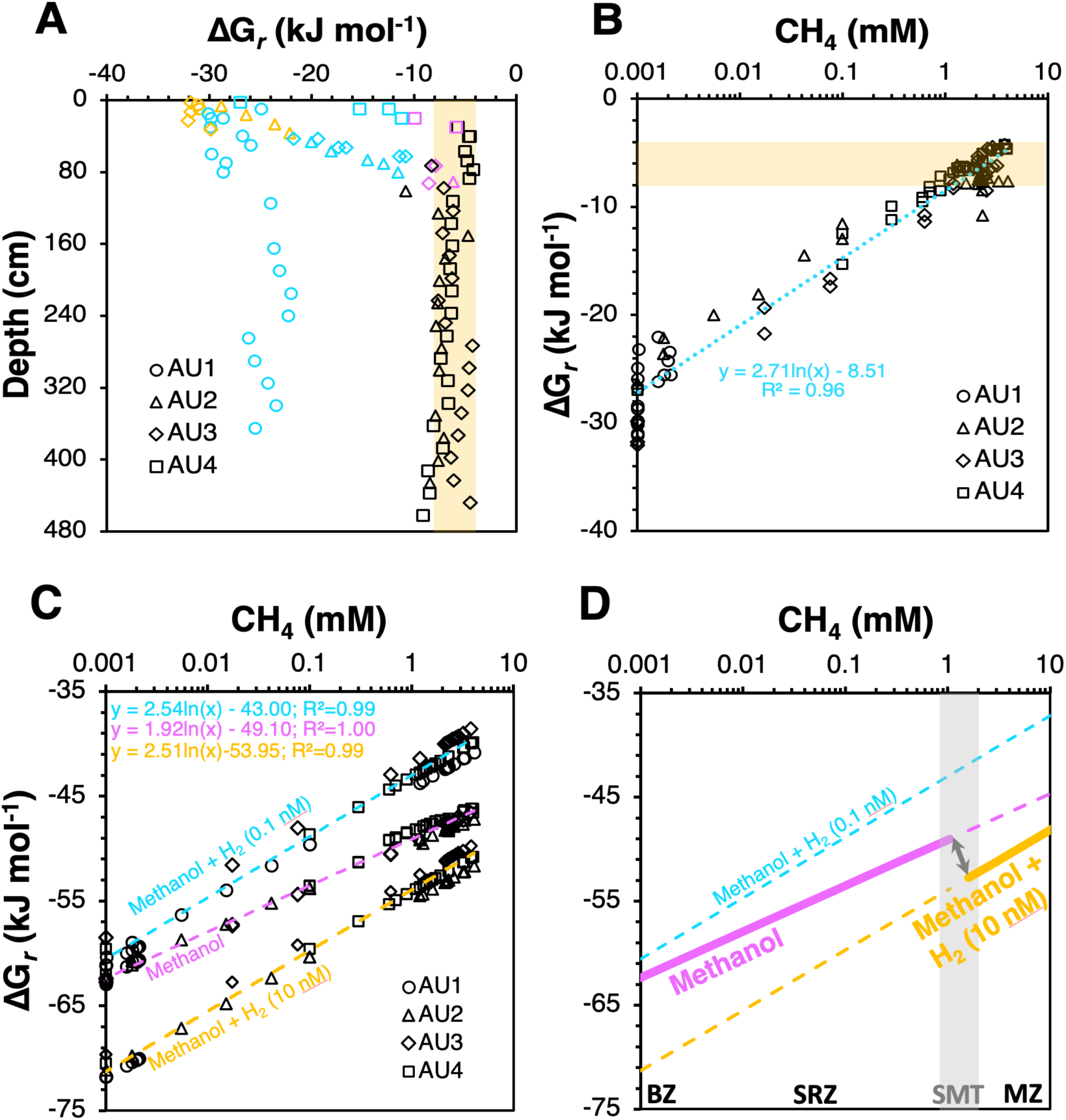
Thermodynamic analyses to investigate energetics of aceticlastic methanogenesis (A-B) and methylotrophic methanogenesis involving methanol (C-D). **(A)** Downcore thermodynamic profile of Gibbs energies of aceticlastic methanogenesis, indicating that free energy yields are higher (Gibbs energies more negative) in bioturbation zones (BTZ; orange symbols) and sulfate reduction zones (SRZ; turquoise symbols) than in sulfate-methane transitions (SMT; magenta symbols) and methanogenesis zones (MGZ; black symbols). **(B)** Relationship between methane concentrations and Gibbs energies of aceticlastic methanogenesis, illustrating the potential for high methane concentrations to present a thermodynamic barrier to this reaction in the MGZ (and SMT). **(C)** Gibbs energies of methanogenesis from methanol and methanol+H_2_ assuming [methanol]=1 nM and two H_2_ concentrations ([H_2_]=0.1 nM; [H_2_]=10 nM) in relation to measured methane concentrations. **(D)** Proposed scenario for a switch in dominant methylotrophic pathway from methanol dismutation in the BTZ and SRZ to methanol reduction in the MGZ that is driven by an increase in H_2_ within the MGZ.

We observe similar overall trends, i.e. higher free energy yields in BTZs and SRZ, and lower free energy yields in SMTs and MGZs, for aceticlastic methanogenesis. Acetate is often considered the most important energy substrate of sulfate-reducing bacteria in marine sediments (e.g., Parkes et al., 1989; Sørensen et al., 1981), and it is generally believed that sulfate reducers outcompete methanogens for acetate (King et al., 1984, Shaw et al., 1984; though also see Sela-Adler et al., 2017). Yet, calculated Gibbs energies of -30 to -10 kJ mol^-1^ reaction in SRZs suggest that even in sulfate-rich sediments, aceticlastic methanogenesis is energetically feasible, with -perhaps surprisingly -higher free energy gains than in underlying MGZs (**Figure 8A**). While the measured, micromolar acetate concentrations are too low for *Methanosarcina* based on previous physiological studies, due to the low-affinity acetate uptake system of the latter (Berger et al., 2012; Jetten et al., 1991), *Methanothrix*, which are known to thrive at acetate concentrations in the low micromolar range, occurred mainly in the SMTs and MGZs (**Figure 3**). So why is aceticlastic methanogenesis not more strongly represented in the BTZs and SRZs given that the reaction is thermodynamically favorable? A potential explanation is the even higher Gibbs energy associate with acetate oxidation coupled to sulfate reduction. While this extra energy may not be conserved via production of higher ATP yields, the resultingly higher thermodynamic drive is likely to benefit sulfate reducers via faster reaction rates. Things change in the underlying sulfate-depleted layers, where aceticlastic methanogens increase in numbers, and are apparently able to drive acetate concentrations down to levels where Gibbs energies are barely exergonic (mostly -8 to -6 kJ mol^-1^; **Figure 8A**). These values suggest that aceticlastic methanogens in MGZs of AU2-AU4 are operating at the absolute thermodynamic minimum, and effectively control porewater acetate concentrations. Anaerobic acetate oxidation (reverse homoacetogenesis), which occurs with methanogenesis in some methanogenic environments (Dyksma et al., 2020; Hattori, 2008), could in theory also control acetate concentrations. Yet, our thermodynamic calculations suggest that at H_2_ concentrations of 10 nM, this reaction is endergonic (**Figure 8B**).

With respect to the observed shift from methyl-dismutation to methyl reduction as the dominant methylotrophic reaction from sulfate-rich to sulfate-dependent sediments, we explore the potential for *in situ* H_2_ concentrations to be the underlying driver (**Figure 8C**). This is based on the previous observation that H_2_ concentrations increase from metal and sulfate reduction-dominated sediments to methanogenic sediments (Hoehler et al., 1998; Lovley & Goodwin, 1988). Assuming methanol as the substrate and 1 µM methanol concentrations, both reactions are always highly exergonic in the sediments studied (-73 to -40 kJ mol^-1^ methanol). Yet, which of the two reactions has higher Gibbs energies depends clearly on H_2_ concentrations. Under “methanogenic H_2_ concentrations” (10 nM), methyl reduction always has higher free energy yields (more negative Gibbs energies) than methyl dismutation. The opposite is the case at low H_2_ concentrations (0.1 nM), as are typical for sulfate and metal oxide reducing sediments. Even if H_2_ concentrations are at 1 nM, where Gibbs energy values are halfway between those for 10 nM and 0.1 nM on the x-axis of **Figure 8C**, methyl dismutation yields higher free energy gains than methyl reduction. Our results offer an energetic explanation for the widely observed dominance of methyl dismutating methanogens as the main methylotrophic methanogens in sulfate-rich surface sediments. By the same token, shifts to methyl reduction as the methylotrophic dominant pathway in deeper layers, which to our knowledge have never been documented, can be expected to be more widespread than currently known and be driven by higher H_2_ concentrations in deeper layers (for conceptual diagram, see **Figure S2**).

### Conclusions

By systematically examining the methane-cycling archaeal community and pathways from coastal eutrophic to off-shore oligotrophic sites, our study shows that the methane-cycling archaeal communities in the continental margin sediments generally contains two complementary “axes of variation”. The first axis reflects the depth-dependent variations that follow the vertical geochemical zonation (i.e., the BTZ, SRZ, SMT, and MGZ), while the second axis captures the site-specific variations. Both axes of variation are largely controlled by the differences of organic matter reactivity and content, which has a major impact on the sulfate penetration depth in the sediment, and the bioturbation activity between sites. By further analyzing the absolute abundances of different methane-cycling archaeal groups, we propose a new framework to classify the methane-cycling archaea in continental margin sediments into ‘surface taxa’, ‘SMT taxa’, ‘MGZ taxa’, and ‘ubiquitous rare taxa’. On one hand this framework is consistent with the existing physiological knowledge, with the maximum abundance of known methyl-dismutating taxa within the *Methanosarcinaceae* in the bioturbation and sulfate reduction zones, and of known methane-oxidizing taxa (ANME-2a-b, ANME-2c, ANME-1a-b) in the sulfate-methane transitions. On the other hand, it indicates that several physiologically uncharacterized groups, including ANME-1d and several new genus-level groups of putatively CO_2_-reducing *Methanomicrobiaceae* and methyl-reducing *Methanomassiliicoccales*, increase significantly in mcrA copy numbers with depth, suggesting the thriving of these groups and their methanogenic lifestyle that is supported by geochemical evidences and thermodynamic calculations. Our study highlights the need for more cultivation research on marine methanogens, given the observed dominance of uncultured groups in methanogenic continental margin sediments, which represent a significant fraction of methanogenic marine sediments.

## Acknowledgments

This study was funded by Swiss National Science Foundation Project 205321_163371 (to M.A.L.). L.D. was sponsored by the Shanghai Pujiang Program (22PJ1404800). The Aurora expedition was jointly funded by the Danish National Research Foundation and European Research Council Advanced Grant (294200, MICROENERGY) (to B.B.J.). We thank the captain, crew, and scientific party of the expedition, as well as Laura Piepgras (MPI Bremen) and Kasper Urup Kjeldsen (Aarhus University) for sampling assistance, and Jean-Claude Walser (ETH Zürich) for bioinformatic assistance.

