## Supplementary materials for "Drivers of methane-cycling archaeal abundances, community structure, and catabolic pathways in continental margin sediments"

**Supplementary tables and figures:**

**Table S1.** Thermodynamic data of aqueous educts and products under standard conditions.

| **Chemical species** | **∆G*_f_*^°^**  **(kJ mol^-1^)** | **∆H*_f_*°**  **(kJ mol^-1^)** | **∆V*_f_*°**  **(cm^3^ mol^-1^)** | **Source** |
| --- | --- | --- | --- | --- |
| H^+^ | 0.0 | 0.0 | 0.0 | Shock et al. (1997) |
| H_2_ | 17.6 | -4.2 | 25.2 | Wagman et al. (1982), Shock and Helgeson (1990) |
| water | -237.2 | -285.8 | 18.0 | Amend and Shock (2001) |
| bicarbonate | -586.9 | -692.0 | 24.6 | Wagman et al. (1982), Shock et al. (1997) |
| formate | -351.0 | -425.7 | 26.2 | Shock and Helgeson (1990) |
| acetate | -369.4 | -486.4 | 40.5 | Shock and Helgeson (1990) |
| methanol | -175.4 | -246.5 | 38.2 | Shock and Helgeson (1990) |
| sulfate | -744.96 | -910.21 | 13.88 | Shock et al. (1997) |
| sulfide | 11.97 | -16.12 | 20.65 | Shock et al. (1997) |

**Figure S1.** Correlation heatmap to examine the relationship between the absolute gene abundances of dominant *mcr*A genus and environmental variables, as well as the correlations between different *mcr*A groups.

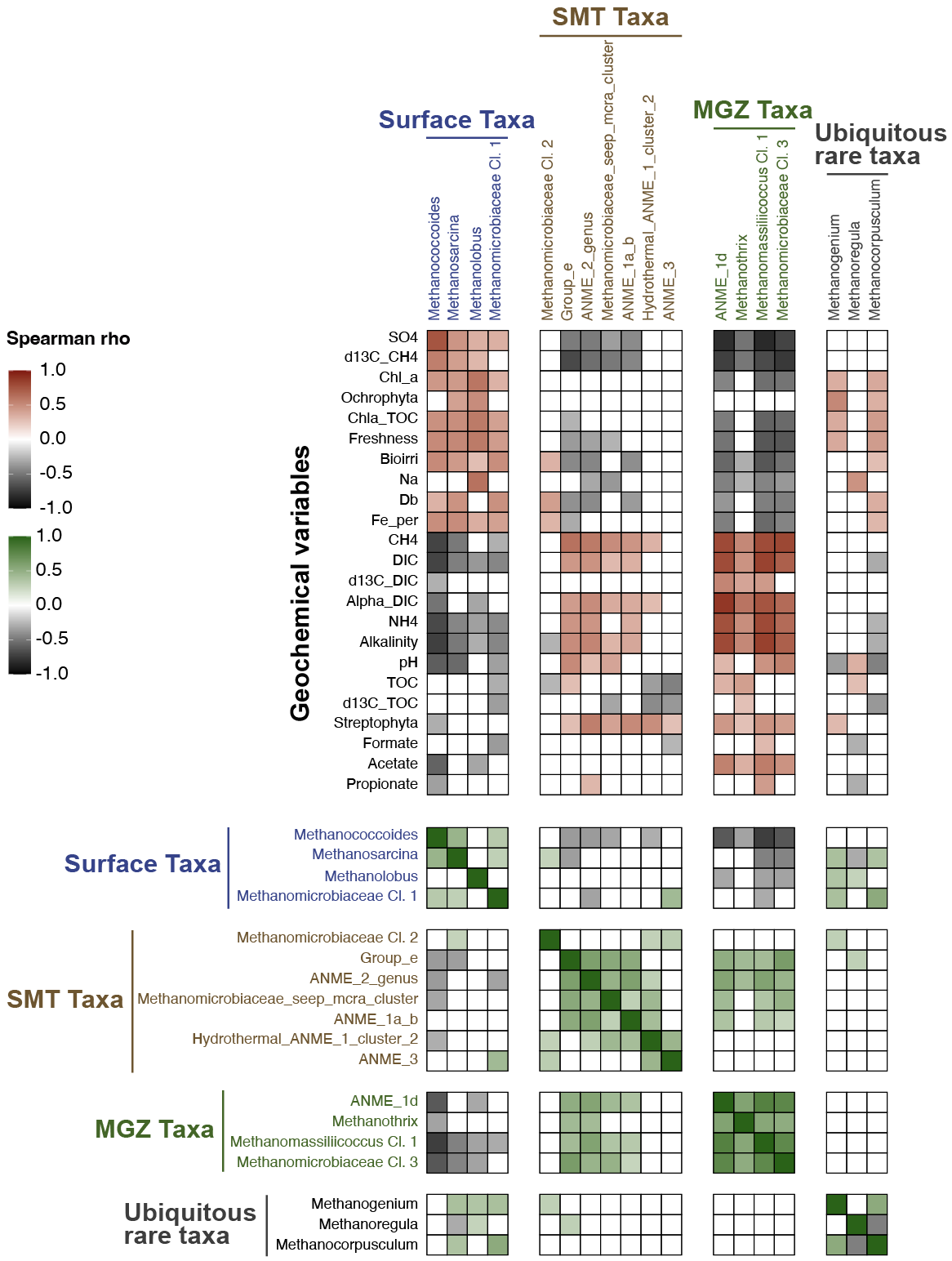

**Figure S2.** Thermodynamic analyses to investigate the energetics of methane-cycling activities under different scenarios.

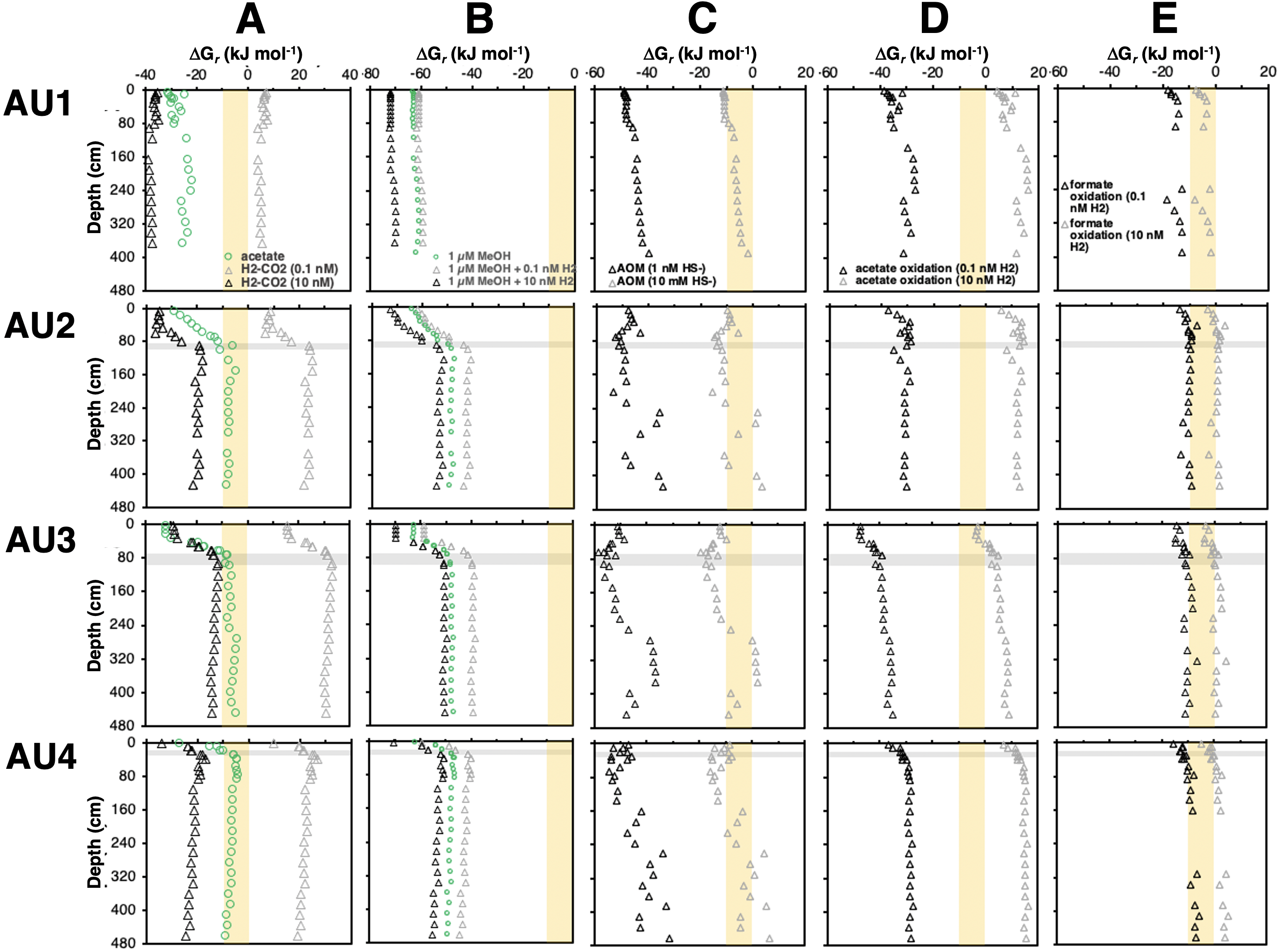
